# The performance of genetic-constraint metrics varies significantly across the human noncoding genome

**DOI:** 10.64898/2026.01.28.701168

**Authors:** Peter McHale, Michael E. Goldberg, Aaron R. Quinlan

**Affiliations:** Department of Human Genetics and Utah Center for Genetic Discovery, University of Utah, Salt Lake City, UT 84112, USA

## Abstract

A longstanding goal in human genetics is to prioritize noncoding loci that, when disrupted, lead to developmental disorders and other Mendelian traits. In pursuit of this goal, multiple metrics have been developed to distinguish neutrally evolving sequences from those subjected to purifying selection. These metrics are commonly evaluated genome-wide, e.g., by computing a precision-recall curve on windows tiling the entire noncoding genome. Here, we identify parts of the noncoding genome where these metrics are distorted due to “bias” in the underlying models of neutral genetic variation and underperform due to a low “signal-to-noise ratio” in the genetic data. The most extreme effects are found for Gnocchi (Chen et al. 2024), the performance of which declines as GC content increases. We trace the source of the bias in Gnocchi to a mismatch between the training set of *de novo* mutations (DNMs) used to adjust expected SNP counts and the genomic windows on which those expected counts are computed. Resolving that mismatch produces a more calibrated version of Gnocchi, significantly improving its interpretability in applications. Looking forward, we suggest annotating constraint scores of noncoding genomic intervals with robust measures of the bias of the corresponding model, allowing users to gauge confidence in those scores.

## Introduction

Computational approaches designed to predict noncoding variant pathogenicity must be improved if they are to have a meaningful impact upon clinical research (Liu et al. 2019). Many of these approaches are based on a simple, but powerful, idea: over the hundreds of millions of years separating species, a single base is likely to have mutated, so the only plausible way to explain why a genomic site is identical between a human and a fish, say, is that whatever mutations occurred there were harmful and eliminated by selection. This deep-time filter is so rigorous that it can detect even weakly constrained elements. Methods like phyloP (Pollard et al. 2010) and GERP (Davydov et al. 2010; Cooper et al. 2005) formalize this idea in a number of ways. (1) They go beyond the vague assertion that a base is “likely to have mutated” by quantifying the expected (neutral) rate of nucleotide substitution between species. (2) They extend the analysis from the comparison of two species to the entire tree of life by accounting for the fact that conservation between, e.g., human and chimp (closely related) is weaker evidence of selection than conservation between human and fish (distantly related). (3) They go beyond a simple binary classification of conservation to answer the question “Is the substitution pattern we see at this site extremely unlikely to have occurred under the neutral model?”

Selection signals derived from sequence conservation are extraordinarily strong and unambiguous but have an Achilles heel: they cannot find genomic elements that are newly constrained in humans, limiting their ability to correctly prioritize noncoding variants. If an element (like an enhancer) becomes functional in the human lineage after the split from chimpanzees, it will not be conserved in chimps, mice, or fish. In scenarios such as these, conservation methods are underpowered (though they can capture elements that have gained function in closely related species, e.g., primates (Boffelli et al. 2003)). The blind spot of conservation-based methods is not small. Rands et al. estimated that only about 25% of the elements under purifying selection in humans are also constrained in mouse (Rands et al. 2014), while, more recently, Huber et al. estimated that most of the noncoding sequence that is functional in human has experienced changes in selection coefficients throughout mammalian evolution (Huber et al. 2020). This process, called “turnover”, had long been suspected from the observation that, as the divergence between mammalian species increases, the predicted amount of pairwise shared functional sequence decreases dramatically (N. G. C. Smith et al. 2004; Meader et al. 2010). Furthermore, most transcription-factor binding events are species-specific, indicative of gain and loss of binding along each lineage (reviewed in (Ponting and Hardison 2011)).

The only way to discover human-specific genetic constraints in the noncoding genome is by using population-based methods. Pleasingly, early deployment of these methods discovered that regulatory elements located near neural genes gained function in the human lineage (Ward and Kellis 2012; Schrider and Kern 2015), and we expect these methods to become increasingly sensitive for the discovery of gain- and loss-of-function discovery as more human genomes are sequenced.

These approaches start with the construction of a model of neutral genetic diversity (e.g., number of SNVs in each of a set of 1kb genomic windows) observed in a healthy human cohort, using features that are known to co-vary with SNV count. Of these, the most well-known feature is the local sequence composition of a given site, whose probability of being polymorphic can increase >10-fold if the sequence is a CpG dinucleotide (Sved and Bird 1990; Bulmer 1986; Schraiber et al. 2024). Beyond local sequence context, the neutral model can include regional features that also covary with SNV density, such as GC content. Chen et al. (Chen et al. 2024) combine the local and regional features in two steps. They first assign each possible SNV a probability of being observed, based on its trinucleotide context and, for CpG transitions, the site’s methylation level. They then adjust the resulting expected SNV count in each window by a factor that accounts for regional features. That factor is estimated separately for each trinucleotide context by modelling whether a site carries a DNM as a function of a panel of 13 candidate regional features, including GC content, each measured at scales from 1 kb to 1 Mb.

With a model of this type in hand, one may identify putative signals of negative selection by searching for anomalous regions in the genome where the observed diversity is significantly lower than the neutral model predicts (Chen et al. 2024; Halldorsson et al. 2022; di Iulio et al. 2018). Alternatively, one may construct a model of *all* genetic diversity, including observed diversity impacted by negative selection. In this approach, the model includes a selection parameter, the value of which is estimated for each genomic window (Dukler et al. 2022). The two approaches are intimately related: as Dukler et al. show, the maximum-likelihood estimate of the selection coefficient in the second approach is the ratio of the observed diversity and that expected under neutrality, the comparison of which is the basis of the first approach.

The success of either approach depends upon how well the underlying model of neutrality tracks the observed SNV density across the noncoding genome: if one inadequately models the neutral mechanisms that, like purifying selection, deplete SNV density, the result is a “biased” model whose predictions of genetic constraint will be flawed. Even if a neutral model can perfectly recapitulate the expected distribution of SNV counts in a locus, one’s ability to predict that the locus is under purifying selection is still limited by how large the “signal” (the depletion of SNV counts due to purifying selection) is relative to “noise” (unavoidable statistical fluctuations in observed SNV counts). Motivated by these concerns, we investigate how existing models’ ability to predict genetic constraint is impacted by variations in model bias and signal-to-noise ratio.

## Results

Consider the Chen model of neutral genetic diversity in humans (Chen et al. 2024), which predicts the distribution of the number of SNVs in a given genomic window, aggregated over 76,156 humans, as a function of local sequence context, nucleotide-level DNA methylation, and a comprehensive set of regional genomic features, including GC content and recombination rates. This model yields a score called Gnocchi, denoted below, which evaluates how typical or unusual an observed SNV count is in the context of the window containing the SNVs,

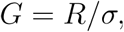

where the residual, *R*, is defined by

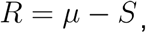

and *S* is the observed SNV count, and *µ* and *σ* are the mean and standard deviation, respectively, of the predicted distribution of SNV counts for that window. A value of Gnocchi near zero indicates that the observed SNV count is close to what one would expect from the neutral model, *µ*. Gnocchi is a *standardized residual*: the residual,*R*, is reported relative to the expected background noise, *σ*. Accordingly,*G≫*1 indicates the SNV *S* count is not only smaller than *µ*, but also smaller than expected statistical fluctuations below *µ*, raising the possibility (but not guaranteeing) that negative selection is operative in that window.

Though Gnocchi is often thought of as a predictor of genetic constraint, it is also a standardized residual of a model of neutral diversity, and, as such, can be used to assess the fidelity of the model. For windows evolving neutrally, a model that matches the truth yields residuals centered on zero at every value of every feature, whatever the true relationship between those features and neutral diversity. Conversely, a systematic departure of the residuals from zero (“bias”) as a function of some feature is evidence that a model’s expectation departs from the truth, whatever it is. The only functional form the residual diagnostic involves is the model’s own; the true functional relationship between genomic features and neutral variation need not be known.

To build intuition for how a discrepancy between a fitted model and the truth appears in standardized residuals, we developed a simulation consisting of the following sequence of steps (Methods). (i) We simulated mutation by random generation of SNVs in a set of mock genomic windows as a function of a mock genomic feature (an example of a real “genomic feature” is “nucleotide content”) distributed normally over windows (**Figure 1A**). The “generative process” is characterized by a “true rate” at which mutations are generated as a function of the genomic feature (black line in **Figure 1B-D**), and sampling from a Poisson distribution with that rate (grey dots in **Figure 1B-D**). (ii) We simulated the development of the model of neutral diversity underlying Gnocchi by estimating the parameters of various models that map the feature value of each window to its SNV count. Like the generative process, the estimated model has its own rate, which we call the “learned rate” (orange line in **Figure 1B-D**) (iii) We simulated Gnocchi by combining the results of (i) and (ii) to compute a standardized residual (grey dots in **Figure 1E-G**).

**Figure 1.**
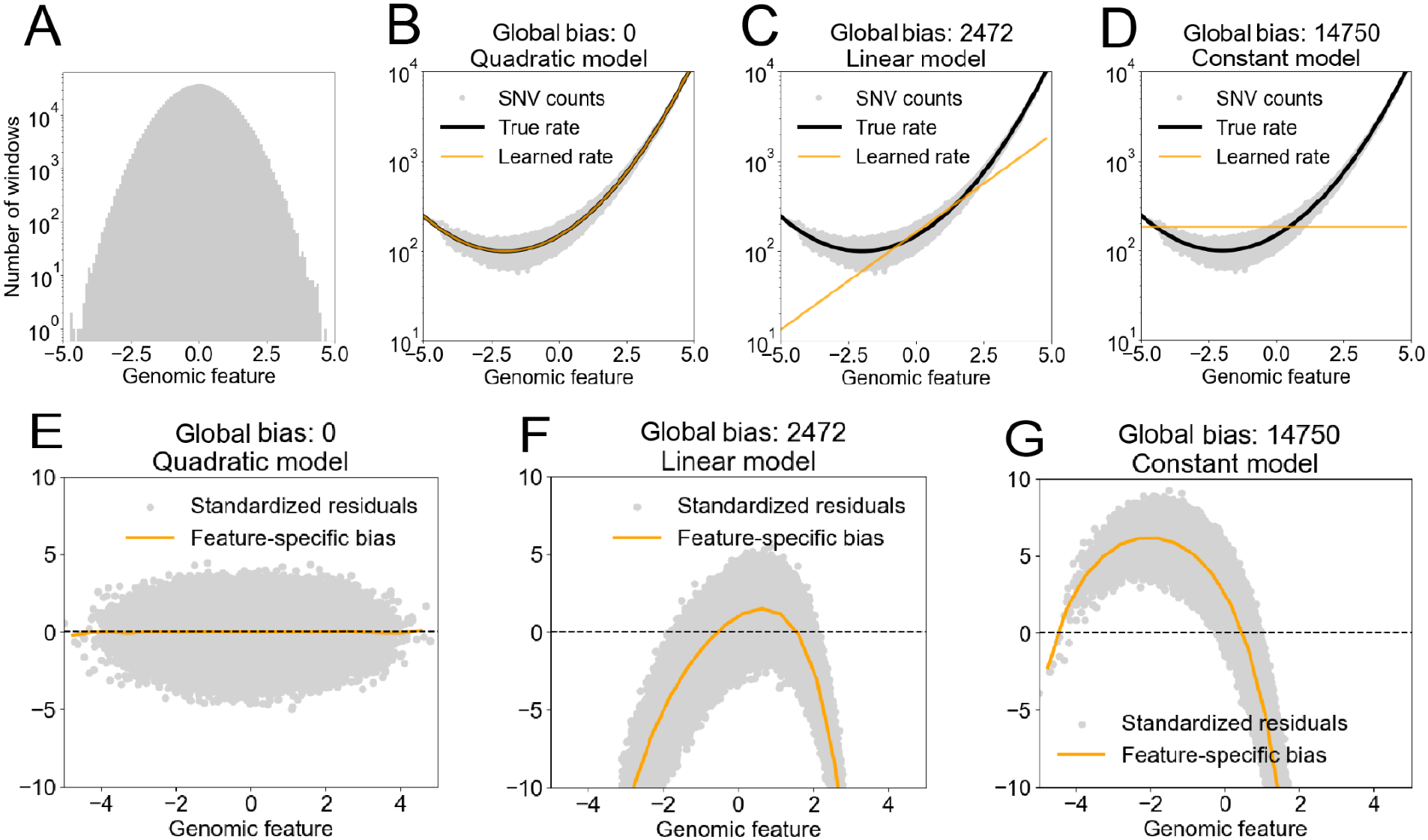
Biased models over- or under-estimate SNV counts, on average, in the tails of the feature distribution. (A) Distribution of a mock genomic feature over simulated genomic windows. (B–D) SNV count of each window (grey) against its feature value, with the true rate from which the counts were drawn (black) and the rate learned by each of three models (orange): quadratic (B), which has the same form as the true rate, linear (C) and constant (D). (E–G) Standardized residuals of each model (grey) and their mean at each feature value, the feature-specific bias (orange); dashed line, zero.

We elected to make the generative process quadratic, not because we expect real SNV counts (or, more precisely, their logarithm) to be quadratic in any genomic feature, but because it is the simplest process that admits two progressively simpler models: one that represents the feature too simply (linear; **Figure 1C, F**) and one that omits it altogether (constant; **Figure 1D, G**). The behavior of the residuals does not depend on the quadratic form, it depends on the deviation of the fitted model’s learned rate from the true rate. The difference between the fitted model and the generative process is quantified by “global model bias”, defined as the squared difference between the learned rate and the true rate, averaged over all windows (Bishop 2006).

We started by using the generated SNVs to estimate a model of the same functional form as the generative process (the “Quadratic model”; **Figure 1B**). Consequently, after training, the estimated model recapitulated the simulation’s generative process, resulting in zero global model bias (**Figure 1B)**. Corresponding to this lack of bias, the distribution of the standardized residual is randomly scattered around zero independent of genomic feature, indicating that observed and predicted SNV counts agree, on average (**Figure 1E**). If the estimated model has a simpler functional form than the generative process, such as when the estimated model is linear but the generative process is quadratic, then the global bias becomes non-zero (**Figure 1C**) because the fit favors the densely populated centre of the feature distribution over its tails (Methods). Associated with the global bias is a “feature-specific bias”, defined as the mean of the residuals as a function of feature value (orange line in **Figure 1F**). The feature-specific bias shifts away from zero, particularly in the tails of the distribution of feature values, indicating that the model systematically over- or under-estimates observed genetic diversity. When the model omits the feature altogether, the global bias is even larger (**Figure 1D**), and the feature-specific bias deviates significantly from zero even in the center of the feature distribution (**Figure 1G**).

Motivated by these results, we sought to visualize the bias of the Chen model by plotting the distribution of Gnocchi on a set of 693,270 *putatively neutral* 1kb intervals (Methods) as a function of three features (**Figure 2A-C**): a measure of nucleotide composition (GC content); a measure of background selection (BGS) computed by (Murphy et al. 2023); and a measure of GC-biased gene conversion (gBGC) computed by (Glémin et al. 2015). This analysis assumes no functional form for the true relationship between these features and neutral diversity; it tests only whether the Chen model’s expectation matches the data. We chose GC content as single-nucleotide substitution probabilities have long been known to decline as GC content increases (Fryxell and Moon 2005; Arndt et al. 2005; Mugal and Ellegren 2011). Additionally, there is evidence that GC content at kilobase scale affects the probability of a CpG site mutating, as the deamination process requires local DNA melting, which in turn requires more thermodynamic energy for bonds between G and C nucleotides (Elango et al. 2008). Finally, even the GC content of the local sequence context of a CpG site may correlate with its mutation rate (Chandra and Gao 2025).

**Figure 2.**
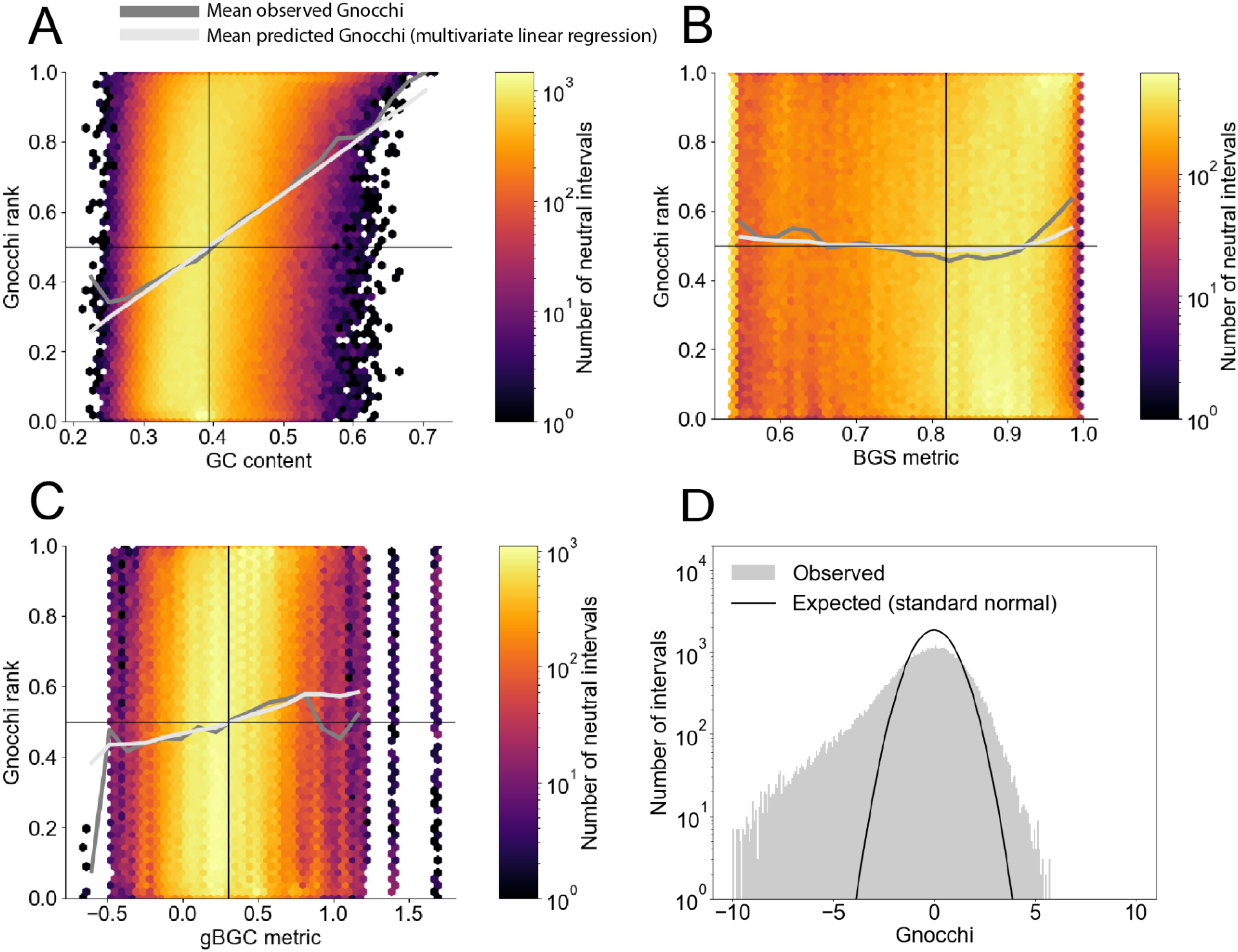
Visualizing the bias of the Chen model of neutral genetic diversity. (A–C) Gnocchi rank against GC content (A), background selection (BGS) metric (B) and GC-biased gene conversion (gBGC) metric (C) in 693,270 putatively neutral 1 kb windows. Gnocchi rank lies between 0 and 1 and averages 0.5 over all windows (horizontal line); an unbiased model would give a mean rank of 0.5 at every feature value. Dark-grey line, mean Gnocchi rank in each feature bin; light-grey line, mean rank predicted by a linear regression on all three features (Table 1); vertical line, feature mean. (D) Distribution of Gnocchi in windows with GC content close to the mean (grey), and the standard normal distribution expected if the model captured all neutral variation (black; Methods).

Though BGS and gBGC metrics are not explicitly included in the Chen model, they are neutral mechanisms well-known to affect diversity in the human genome (McVicker et al. 2009; Pouyet et al. 2018; Murphy et al. 2023), and have been modeled when inferring natural selection in the context of population genetics, e.g., (Telis et al. 2020). BGS, as a process, reduces variation in regions linked to sequences constrained by purifying selection (for a review see (Charlesworth 2013)). GC-biased gene conversion (reviewed in (Duret and Galtier 2009)) describes how mismatch repair of heteroduplexes formed during meiosis is biased towards GC alleles, and thus against AT alleles, altering their probability of fixation. Therefore allele frequency is also altered in a manner that mimics natural selection though is neutral (Duret and Galtier 2009). As such, like GC content, both BGS and gBGC are neutral processes that lead to reduced genetic diversity which could masquerade as selection to models that fail to account for them.

We observe bias (represented by the deviation of the dark-grey line from *y* = 0.5 in **Figure 2A-C**) in the Chen model in rare windows that harbor extreme levels of background selection, or GC-biased gene conversion, but particularly those windows with extreme levels of GC content. Note, however, that the lack of bias in the Chen model at intermediate feature levels does not imply that the model cannot be improved there. In fact, we observe that the variance of Gnocchi in that vicinity is larger than what would be expected (**Figure 2D**), implying that there remains variation in SNV count that the Chen model is unable to capture even at genomic windows whose feature values are observed frequently across the genome.

We next sought to measure how each feature *independently* contributed to the Chen model’s bias. The slopes of the dark-grey lines in **Figure 2A-C** do not measure these contributions.

Instead, these slopes represent not only the direct effect of any given feature (e.g, GC content) on model bias, but also the indirect effect of the complementary features (BGS metric and gBGC) due to correlation among these features. To capture the independent effect of each feature on model bias, we fit a linear model of all features to Gnocchi (Methods). The estimated parameters of the linear model represent the direct effects of the corresponding feature on the bias of the Chen model (**Table 1**). We find, for example, that the direct effect of the BGS metric on Gnocchi is negative, implying that stronger background selection (smaller values of the BGS metric) is associated with larger values of Gnocchi. This accords with our naive expectation that a model of neutral diversity that does not explicitly account for BGS (as is the case with the Chen model) ought to overestimate diversity where background selection is strong (assuming that BGS does not significantly co-vary with features that the Chen model does include).

**Table 1.**
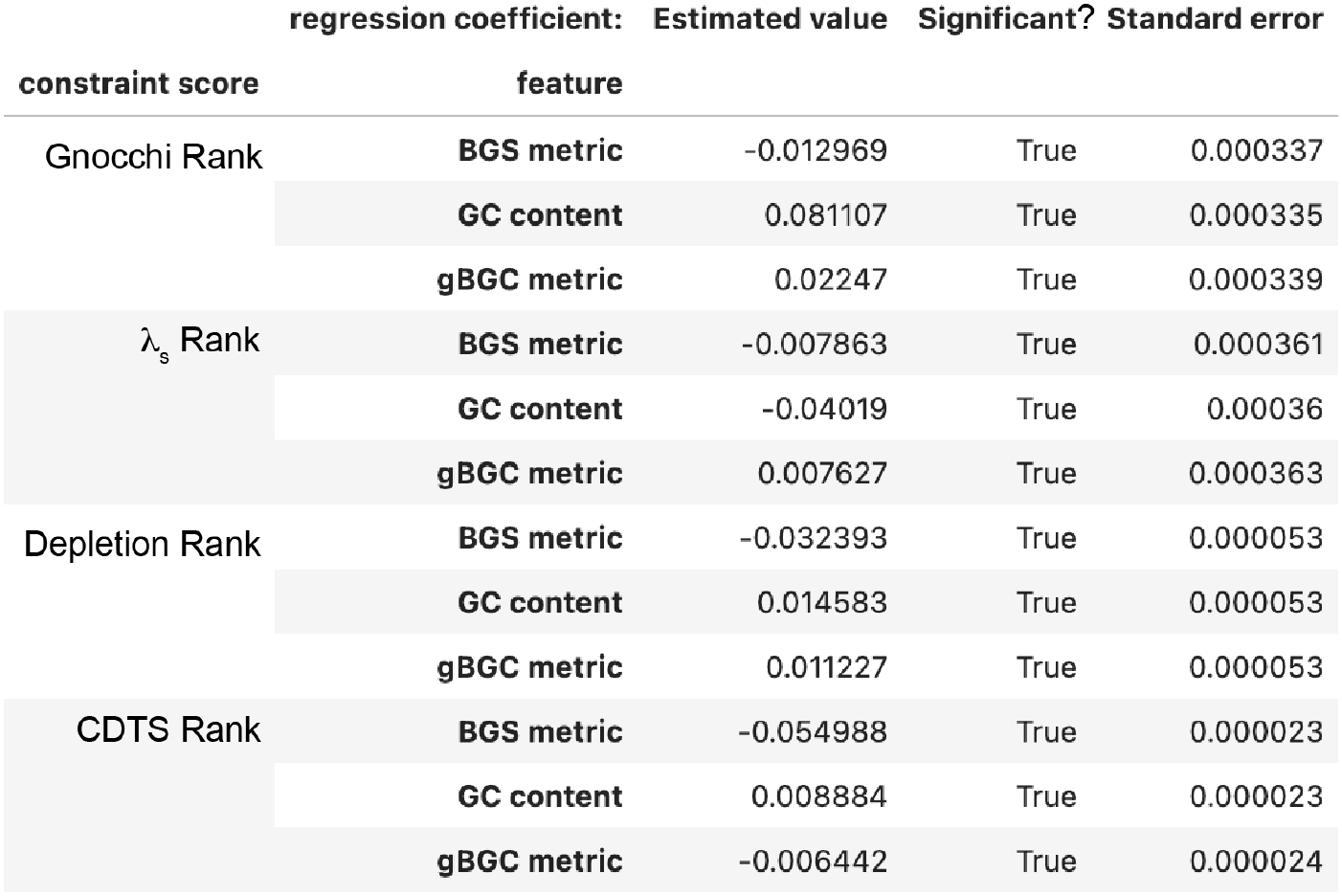
The direct effect of each feature on the bias of the neutral models underlying each constraint metric. The regression coefficients may be compared not only within a given model, but also among models, since the features are standardized prior to fitting, and the target variables (ranks) in the fitting process are uniformly distributed between 0 and 1.

Interestingly, though a positive value of Gnocchi *erroneously* predicts constraint in the neutral windows subject to strong BGS, it *correctly* predicts constraint nearby, by definition of BGS. Since we standardized all features prior to fitting our linear model to Gnocchi, we may compare the values of the corresponding regression coefficients, revealing that GC content has the largest effect (in magnitude) on model bias: GC content explains almost as much variance in Gnocchi as explained by the three features jointly (compare for one feature vs three features, respectively, in **Table 2**).

**Table 2.** The proportion of variance in each constraint score that is explained, *ρ*^*2*^, by all three features considered in this study, and by the single feature with the largest direct effect on bias.

| **constraint score** | **features** | *ρ*<sup>2</sup> |
|---|---|---|
| Gnocchi Rank | [GC content, BGS metric, gBGC metric] | 0.089556 |
| Gnocchi Rank | [GC content] | 0.082728 |
| λ<sub>s</sub> Rank | [GC content, BGS metric, gBGC metric] | 0.020279 |
| λ<sub>s</sub> Rank | [GC content] | 0.019076 |
| Depletion Rank | [GC content, BGS metric, gBGC metric] | 0.014899 |
| Depletion Rank | [BGS] | 0.010312 |
| CDTS Rank | [GC content, BGS metric, gBGC metric] | 0.038128 |
| CDTS Rank | [BGS] | 0.036908 |

To assess the generality of these results, we repeated our analysis for three other constraint metrics: λ_*S*_ (Dukler et al. 2022), Depletion Rank (Halldorsson et al. 2022) and CDTS (di Iulio et al. 2018) (Methods). The results, presented in **Tables 1** and **2**, and **Supporting Figure 1**, are similar to those obtained for the Chen model in the sense that all models develop biases as one moves into the tails of the feature distributions, though the details differ. For example, whereas high GC content biases Gnocchi above zero (rank > 0.5), it biases λ _*S*_ below zero (rank < 0.5) (**Supporting Figure 1)**. These high-GC-content intervals may be enriched for promoter regions, where the authors of the λ_*S*_ metric observed a similar bias, which they attributed to misspecification of the underlying neutral model there (Dukler et al. 2022). Another difference between the Chen model and the others is that BGS, not GC content, explains most of the bias observed in Depletion Rank and CDTS (e.g., **Table 2**).

The procedure we adopted to identify the feature that drives model bias is closely related to Gradient Boosting (Friedman 2001), a technique for improving a model by regressing its residuals onto its features, and then adding the resulting “residual model” to the predictions of the original model. Viewed this way, the *ρ*^2^ values reported in **Table 2** measure how much an improved modeling of the corresponding features (e.g., via Gradient Boosting) is expected to improve the underlying model of neutral diversity. Comparing the *ρ*^2^ values among features and models leads to two interpretations. First, the largest improvement to the Chen model is expected to come from more faithful representations of sequence composition (for which GC content is a surrogate), whereas the largest improvements to the models underlying Depletion Rank and CDTS are expected to come from better representations of BGS. Second, of all the models, the Chen model stands to benefit most from an improved modeling of the three features considered in this study.

The purpose of constraint metrics, such as Gnocchi, is to discover “genetic constraint”, i.e., to distinguish genomic windows that are neutral from those that are under purifying selection. To understand how well this approach is likely to work under various circumstances, we returned to simulation. The standard approach to modeling purifying selection is to use forward-time population genetic simulations, e.g., via stdpopsim (Adrion et al. 2020; Lauterbur et al. 2023; Gower et al. 2025). Though stdpopsim incorporates a host of neutral processes, each of them is expected to contribute to “noise” (unavoidable statistical fluctuations in observed SNV counts in genomic windows), obscuring the patterns we wish to illustrate with our simulations. To minimize this noise, and therefore maximize our ability to recover “signal” (the depletion of SNV counts due solely to purifying selection), we decided instead to return to our simple simulation of neutral diversity (**Figure 1A-D)**, which is stripped of most of the neutral processes included in tools like stdpopsim. We extended the simple simulation to model purifying selection by simply reducing the SNV counts of a random subset of our simulated windows, labeling them “constrained”, and labeling the rest “neutral”. We then built a constraint predictor by thresholding residuals corresponding to each of the models of neutrality depicted in **Figure 1B-D**: windows with residuals greater than the threshold were predicted to be constrained, whereas those with residuals lower than the threshold were predicted to be neutral. Finally, we assessed the performance of those models by comparing predicted labels with actual labels.

Even if a model is unbiased, its performance is limited by the signal-to-noise ratio (SNR): the ratio of the “signal” we are trying to recover (the depletion of SNV counts due to purifying selection) and the unwanted background “noise” we must contend with (unavoidable statistical fluctuations in observed SNV counts). A biased model is limited further, since errors in its expected SNV counts can mimic, or mask, the depletion caused by purifying selection. To separate the effects of SNR and model bias on performance, we analysed windows within “strips” of feature values (**Figure 3A–C**). Within a narrow strip the feature value is effectively constant, and so therefore is each model’s prediction, so all models perform alike and performance is limited by SNR alone (Methods). Across a wide strip, however, the feature varies enough for each model’s bias to affect performance as well.

**Figure 3.**
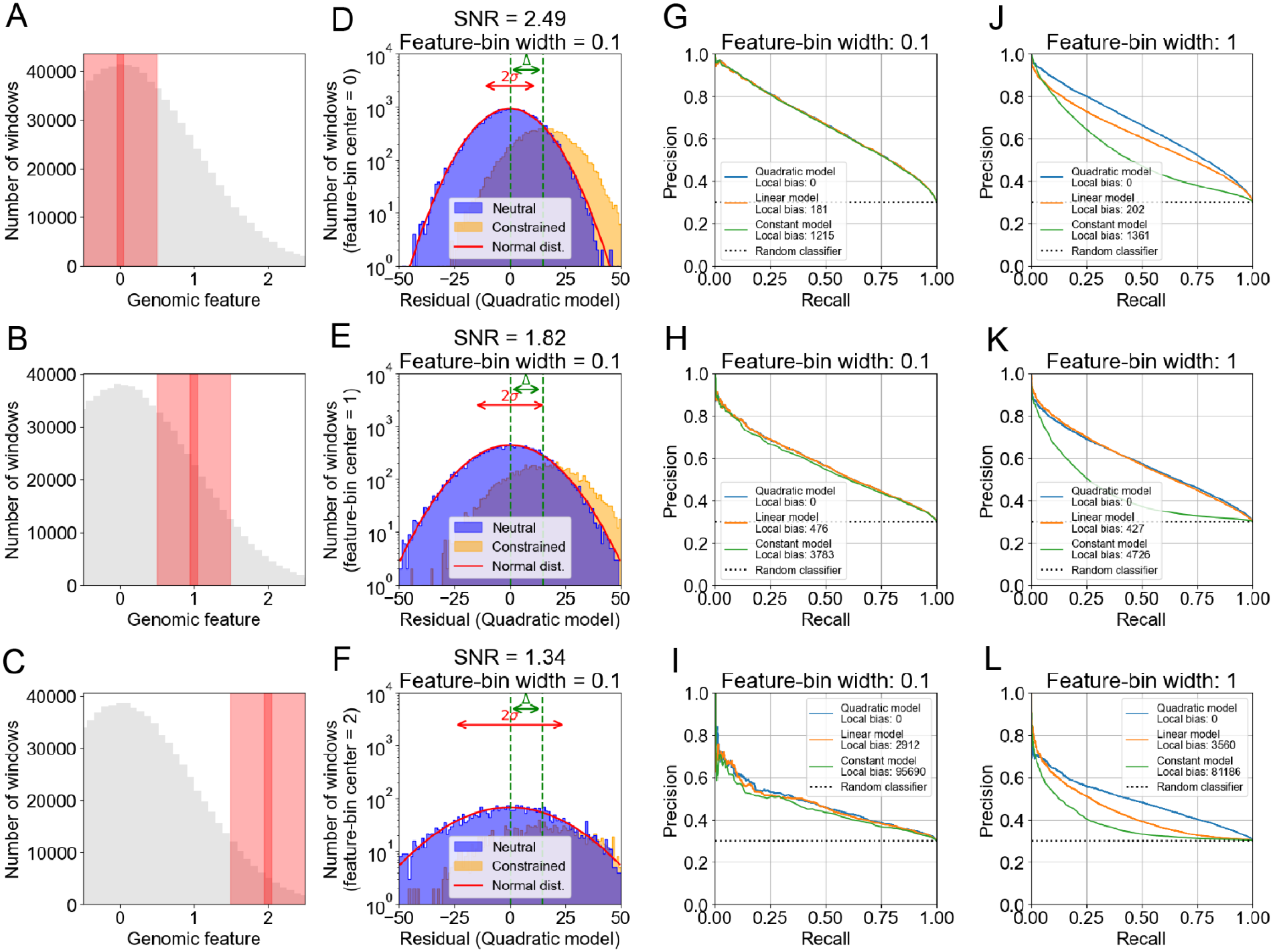
SNR and model bias are both expected to significantly affect the ability to call constraint. Simulated windows as in Figure 1, of which a random subset was labelled constrained and had its SNV counts reduced (Methods). Each row analyses windows whose feature value lies in a strip centred on 0 (top), 1 (middle) or 2 (bottom). (A–C) Distribution of the feature, with the narrow (width 0.1, dark red) and wide (width 1, light red) strips. (D–F) Residuals under the quadratic model of neutral (blue) and constrained (orange) windows in the narrow strip; red curve, normal distribution for comparison; *σ*, standard deviation of the neutral residuals; Δ, depletion of SNV counts in constrained windows. (G–I) Precision–recall (PR) curves for predicting the constrained windows in the narrow strip from their residuals. (J–L) As G–I, but for the wide strips; each model’s local bias is given in the legend.

We first took narrow strips centred on three different values of the genomic feature (**Figure 3A–C**), and in each case computed the SNR (**Figure 3D-F**; Methods). Regardless of model bias, performance declines (**Figure 3G-I**) as SNR declines (reflected in an increasing overlap of the residual distributions of the neutral and constrained windows; **Figure 3D-F**).

We then widened each strip enough to expose the additional impact of bias on the performance of the three models, yet not enough to alter the SNR in each case (compare narrow with wide red strips in **Figure 3A-C**). With this change, not only does performance decline due to declining SNR as found previously (notice the similarity of the blue lines in **Figure 3J-L** with the corresponding blue lines in **Figure 3G-I**), but also because of increasing bias for any given SNR (for each panel of **Figure 3J-L**, compare the three colored lines).

Given that model bias and SNR are expected to jointly affect the ability to predict constraint in a way that is impossible to predict without knowledge of that which we are seeking (genetic constraint), we set out to empirically measure performance of Gnocchi as a function of GC content, BGS and gBGC. We designed a simple classifier that predicts that a window is constrained if Gnocchi exceeds a threshold. Since the main purpose of, and main challenge facing, scores such as Gnocchi is to identify constraint in the noncoding genome, an example of which may be a conserved enhancer, we constructed a set of noncoding windows and labeled those that overlap a GeneHancer enhancer (Fishilevich et al. 2017) “constrained”, and those that don’t “neutral” (Methods). Measured against this truth set, Gnocchi clearly does better than random guessing (compare black line and dashed line in **Figure 4A)**, but its performance in GC-rich windows is significantly reduced (red lines); conversely, in GC-poor windows, performance is better than expected (blue lines). Since Gnocchi is biased at low GC content (**Figure 2A**), and in light of the results in **Figure 3**, we conjecture that the increased performance at predicting constraint in GC-poor regions is due to an increased SNR. Similar variability in performance was observed as a function of BGS and gBGC (**Supporting Figure 5A, B**). These results illustrate how model bias and/or SNR jointly generate significant variation in ability to discover constraint under the Chen model, depending upon the nature of the windows being interrogated.

**Figure 4.**
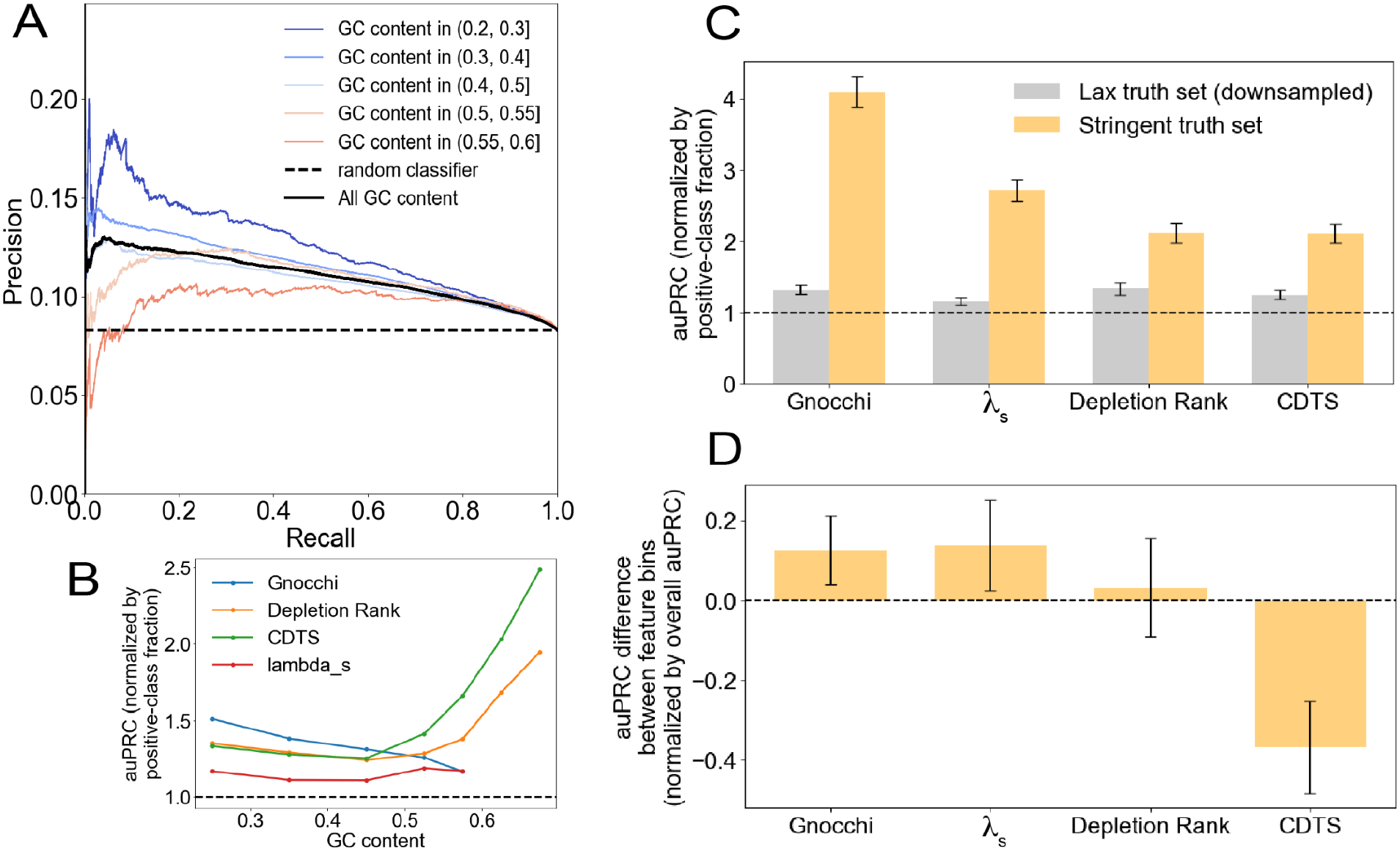
The ability of models of neutral diversity to predict constraint varies with the nature of the genomic windows. (A) Precision–recall curves of Gnocchi on the lax truth set (Methods), over all windows (black) and within each GC-content bin (coloured); dashed line, random classifier (Methods). (B) Normalized auPRC of four constraint metrics on the lax truth set, against GC content. auPRC is the area under the Precision–Recall Curve; normalizing divides it by the fraction of windows labelled constrained, so that a random classifier scores 1 (dashed line). (C) Normalized auPRC of each metric on the stringent truth set (Methods) and on the lax truth set downsampled to the same size. (D) Difference in normalized auPRC between high-GC (0.4–0.7) and low-GC (0.2–0.375) windows, on the stringent truth set. C and D, mean ± s.d. over 1,000 bootstrap samples; dashed lines, random classifier (C) and no difference (D).

To compare constraint metrics, we summarized each precision–recall curve by the area under it (auPRC), divided by the fraction of windows labelled constrained so that a random classifier scores 1 (Methods). Similar trends in performance are observed for all constraint metrics, with the notable exception of CDTS and Depletion Rank, which increase in performance in GC-rich and/or gBGC-rich regions (**Figure 4B**, and **Supporting Figure 5C and D**, respectively).

Interestingly, these data show that, of all the models, the Chen model has the best *average* performance, i.e., when evaluated at typical values of the features (vertical black lines in **Figure 2A-C**), even though it has the greatest bias (see *ρ*^2^ in **Table 2**).

The number of windows in the truth set that we used to measure performance is large, enabling us to detect performance anomalies that occur deep in the tails of the feature distributions. But that truth set is also lax, as not all windows that overlap a Genehancer enhancer are expected to be highly constrained. For example, 18.4% of the noncoding genome is covered by Genehancer enhancers (Martini et al. 2023), but others have estimated that only 4.51% of the noncoding genome is under human-specific selection (Huber et al. 2020). We therefore identified noncoding windows that we conjectured would be under strong negative selection because they regulate the expression of essential genes (Methods). We combined them with an equal number of noncoding windows not overlapping any enhancer to form a small but stringent truth set. We used it to re-assess the performance of the various constraint metrics at predicting constraint. We observed that overall performance increases on moving from the lax to the stringent truth set, particularly for the Gnocchi and λ _*S*_ metrics (**Figure 4C** and **Supporting Figure 5E, F**), providing a post-hoc validation of stringency. Even though the stringent truth set is small–making estimates of performance in the tails of the feature distributions noisy–we were nevertheless able to detect associations between performance and features (**Figure 4D** and **Supporting Figure 5G,H**), as we observed for the lax truth set (**Figure 4B** and **Supporting Figure 5C,D**).

We were curious as to the source of the strong GC-dependent bias we observe in Gnocchi (**Figure 2A**) but not in Depletion Rank (**Supporting Figure 1**). These scores differ in the existence of an adjustment to the expected SNV counts in Gnocchi that is absent in Depletion Rank (Methods). This suggests a hypothesis that the adjustment causes the observed bias in Gnocchi. We tested the hypothesis by removing the adjustment, producing an “ablated” version of Gnocchi. Indeed, this reduced the bias of Gnocchi to levels comparable to Depletion Rank (**Figure 5A**), implying that the adjustment is primarily responsible for Gnocchi’s bias.

**Figure 5.**
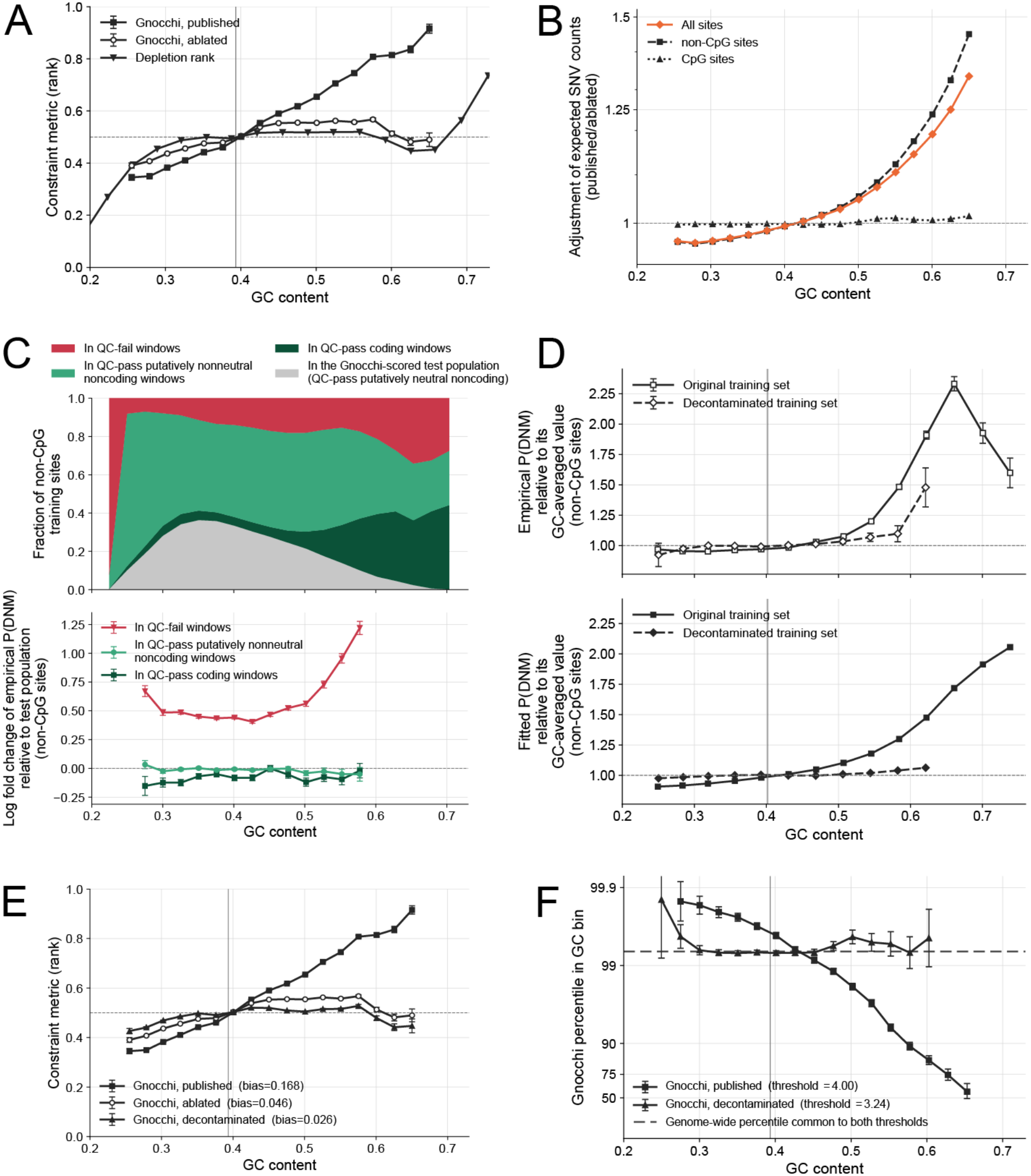
Bias in Gnocchi is driven by bias in a training set underlying the GC-dependent adjustment of expected SNV counts. (A) Mean rank of the constraint score per GC bin for published Gnocchi, ablated Gnocchi and Depletion Rank. (B) Adjustment of expected SNV counts per GC bin at all sites, non-CpG sites and CpG sites. (C) Upper, the fraction of 4,367,225 non-CpG training sites per GC bin lying in each class of 1 kb window; grey, the test population. Lower, the empirical DNM probability in each other class relative to the test population. (D) Empirical (upper) and fitted (lower) DNM probability per site for the original and decontaminated training sets. (E) As A, with decontaminated Gnocchi added. The legend gives each curve’s bias, defined as the average distance of the plotted curve from the rank = 0.5 line. (F) Percentile within each GC bin of E at which each score’s threshold falls; dashed line, the percentile both thresholds share over all windows. Error bars, 95% Wilson intervals. A, E and F are computed on the 693,270 putatively neutral noncoding 1 kb windows of Figure 2, and C and D on non-CpG training sites. Vertical lines mark the mean GC content of those windows (A, E, F) or of the original training sites (D). Error bars, ±1 s.e. unless stated.

The adjustment is designed to control for the known covariation of regional genomic features with the occurrence of SNVs. However, as far as GC content at CpG sites (NCG) is concerned, Chen et al. did not implement the adjustment because GC content is implicitly accounted for in their workflow prior to adjustment. Specifically, they model SNVs as a function of DNA methylation (**Supporting Figure 7A**), which in turn covaries with GC content (**Supporting Figure 7B**). We reasoned that the adjustment of expected SNV counts at the remaining sites — non-CpG sites — must therefore drive Gnocchi’s bias. To test this, we quantified the adjustment as the ratio of expected SNV counts under the published model to that under the ablated model using: (i) all sites; (ii) only CpG sites; and (iii) only non-CpG sites. As expected, the adjustment is dominated by non-CpG sites, with hardly any adjustment made at CpG sites (**Figure 5B**). We conclude that Gnocchi’s bias is driven by the adjustment applied at non-CpG sites exclusively, and therefore consider only non-CpG sites in the subsequent analyses.

Though the adjustment is used to correct the expected counts of SNPs in quality-controlled (QC-pass) neutral windows, Chen et al. elected to compute the adjustment by modeling the occurrence of DNMs, not SNPs. This was done via the estimation of the probability of a DNM occurring at a given site, P(DNM), from a training set of sites at which DNMs have or have not been observed. This raises the question of how representative that training set is of the test set of sites used to compute expected SNP counts, and to subsequently score Gnocchi at the window level. **Figure 5C** (upper) shows that a majority of the sites in the training set are actually not representative of the test set at all, either because they fail to pass QC, sit in putatively nonneutral noncoding regions, or are part of the coding genome. We refer to such sites as “contamination” of the training set. Contamination would be harmless if it didn’t affect P(DNM). Unfortunately, it does. Not only is the empirical P(DNM) at training sites in coding regions reduced because of the action of purifying selection, it is massively inflated in a strongly GC-dependent fashion in training sites that fail QC (**Figure 5C**, lower), possibly because false DNMs are enriched there. Thus the DNM training set that underlies the regional adjustment in Gnocchi is contaminated with consequential sites.

Could contamination of the DNM training set also cause the bias we observe in Gnocchi? To find out, we removed the offending sites from the training set and used Chen et al.’s workflow to recompute Gnocchi. As expected, decontamination reduced the dependence of empirical P(DNM) on GC content (**Figure 5D**, upper), which propagated to corrections of estimated P(DNM) and Gnocchi scores that are much less sensitive to GC content (**Figure 5D**, lower and **Figure 5E**). Not only did decontamination reduce the bias of Gnocchi relative to its published version, it also did so with respect to the ablated version, as one would expect of a properly calibrated adjustment for GC content.

To illustrate the utility of debiasing Gnocchi, consider how it is normally used in practice. Faced with a region of the noncoding genome, an analyst gauges the significance of its Gnocchi score by asking whether it is larger than some threshold that lies in its right tail where ostensibly constrained windows are over-represented. Implicit to this procedure is the assumption that the number of constraint calls a given threshold yields is *independent* of genomic features. For the published version of Gnocchi, that is not true: a given threshold is more strict than average in GC-poor regions, calling relatively fewer regions “constrained”, and considerably less strict than average in GC-rich regions, making many more calls than expected (**Figure 5F**). In contrast, the strictness of our decontaminated version of Gnocchi is largely independent of GC content (**Figure 5F**). Producing a properly calibrated version of Gnocchi is powerful: otherwise an analyst considering a given genomic region needs to tailor his threshold to the peculiar distribution of Gnocchi specific to the GC content of that region. Moreover, it is conceivable that Gnocchi is biased as a function of other genomic features not explored in this manuscript, in which case the analyst would need to condition on those, as-yet-unknown features too before computing his threshold. In short, debiasing makes a given Gnocchi score mean the same thing at every locus in the noncoding genome.

To assess the performance of the decontaminated version of Gnocchi, we returned to the lax truth set considered earlier. Averaged across thresholds, decontaminating Gnocchi has no significant effect on discovery in any GC stratum (**Supporting Figure 8C**), preserving the decline in performance with GC content that we previously observed (**Figure 4B**). That decline must therefore be explained almost entirely by variation in SNR, the only other effect identified in this study that could affect discovery performance. Indeed, this possibility was admitted by our simulations, e.g., any of the operating regimes illustrated in **Figure 3G-I, K**.

When the Gnocchi threshold is fixed, instead of averaged over, decontaminating Gnocchi does have beneficial consequences for the discovery of constraint. Not only does decontaminating redistribute discovery toward sequence where a call is worth more (from GC bins where calls have lower evidential weight to GC bins where calls have more; **Supporting Figure 8A**,**B;** Methods), it also moderately increases the evidential weight of those calls (at matched call budgets; **Supporting Figure 8F, I**).

## Discussion

Sequence conservation leverages the power of deep evolutionary time to provide strong signals of purifying selection, but this approach is limited to the discovery of selection common to a group of species. This forces us to look at human-specific diversity-based methods in order to find human-specific selection. The trade-off is that inference of selection via methods that rely on segregating sites is confounded in a number of ways. (1) Though rare, the fixation of an advantageous allele can drag linked, neutral DNA with it, sweeping away existing variation in that genomic region (J. M. Smith and Haigh 1974; Stephan 2019), and naively leading to false-positive calls of purifying selection. (2) During a population bottleneck, rare alleles tend to be lost quickly, before heterozygosity has been eroded, resulting in a transient imbalance between heterozygosity and allele number (Luikart et al. 1998; Amos and Hoffman 2010) that one could falsely interpret as the active removal of deleterious alleles in that interval by purifying selection, if demography is not accounted for. (3) The demographic history of a population can reduce the ability of natural selection to remove weakly deleterious variants (Lohmueller 2014), and so diversity-based methods, which rely on the removal of deleterious variants, will not discover constrained regions harboring those variants. Here, we identify additional factors that confound our interpretation of diversity-based methods for genetic-constraint calling.

We show that current models of neutral genetic diversity are biased, distorting how they uncover constraint in the human noncoding genome in rare but important contexts, e.g., high GC content (enriched for promoters) or strong background selection (enriched for proximity to genes), depending upon the model. The failure is not that constraint goes undetected in those contexts: at higher GC content Gnocchi actually recalls more windows, not fewer (**Supporting Figure 8A**). It is that a given score ceases to mean the same thing across the genome, so a threshold chosen from a genome-wide distribution of Gnocchi scores selects a far larger and less informative set of windows in higher GC content sequence (**Figure 5F, Supporting Figure 8A**). This mirrors similar findings in the coding genome for a variety of organisms (Yap et al. 2010; Venkat et al. 2018; Ratnakumar et al. 2010).

Similar to the difficulty in capturing edge cases when training machine-learning models to drive cars or answer questions in natural language, models of genetic diversity tend to capture the average behavior seen in a putatively neutral training set, at the expense of incurring large prediction errors on classes of genomic intervals that are rare in that training set. Capturing only average behavior is a blessing when rare events are true anomalies (e.g., constrained windows that are erroneously part of the training set), but a curse when they are not (e.g., truly neutral windows with high GC content).

We find that the model with the highest performance on both the lax and stringent truth set—the Chen model—is also the model with the largest bias. That association across models does not, however, imply a trade-off within one: when the bias is removed from the Chen model itself, average performance does not fall (**Supporting Figure 8C**), but can in fact increase (**Supporting Figure 8I**). Interestingly, the direct effect of GC content on bias is larger than that of gBGC and BGS, even though the former was explicitly included in the model whereas the latter were not. Since performance of the Chen model declines with GC content (**Figure 4A**), and since GC content is positively correlated with Gnocchi (**Supporting Figure 6A**,**B**), Gnocchi is most unreliable precisely where it predicts the most constraint (**Supporting Figure 8A)**.

Moreover, given that GC content is auto correlated across the human genome, forming “isochores” (**Supporting Figure 6B**), we expect these false calls of constraint under the Chen model to cluster in genomic space. The relatively high performance of the Chen model (marginalized over all genomic feature values) may result from how comprehensive its set of regional features is. Its bias, however, arises mostly not from the functional form assumed by those features in the neutral model but from the training population used to estimate their effect on genetic variation. Refitting the same model on the subset of training sites that are actually characteristic of scored noncoding windows removes most of the bias (**Figure 5E**). The residual bias that remains may indicate that the way these features, particularly GC content, are included in the model is still not flexible enough to capture neutral genetic diversity fully.

Deep-learning models, such as those used to predict the density of noncoding SNVs (Fang et al. 2022), or their effects on fitness (Benegas et al. 2024; Trevino et al. 2021), do not suffer from such limitations on model complexity. Nevertheless, a deep-learning model mapping genomic features onto SNV density may still fall foul to the type of biases we illustrated here, as recently illustrated for other genomic prediction tasks (Kathail et al. 2024b; Bajwa et al. 2024). To the extent that this failure mode is due to insufficient drive to improve the fit in sparsely populated parts of feature space, a possible solution is to up-weight training examples there, e.g., by weighting a genomic interval in inverse proportion to the number of intervals similar to it (Morcos et al. 2011).

In the meantime, we recommend annotating each genomic interval not only with its constraint score, but also a measure of how biased the score is expected to be in that interval, as a way to gauge confidence in the score. One approach worth considering is to group genomic intervals by the similarity of their regional features, computing the distribution of constraint scores in each group, and assigning to each window the mean of the constraint-score distribution for the corresponding group.

## Methods

### Simulation of neutral SNV counts and model bias (Figures 1 and 3)

We assigned each of 1,000,000 mock genomic windows a feature value x drawn from a normal distribution, and drew its SNV count from a Poisson distribution whose rate, the *true rate*, is a quadratic function of the feature: λ(x) = exp(A + Bx + Cx^2^). We fit three models by Poisson regression, in which the logarithm of the *learned rate* is a quadratic, linear or constant function of the feature, to the SNV counts. The fits used only the windows labelled “neutral” (see “Simulation of constraint calling” below). Because Poisson regression maximizes the likelihood summed over all windows, a model whose functional form cannot follow the true rate (e.g., linear or constant) is fit most closely where windows are dense, near the centre of the feature distribution, and least closely in the sparse tails. For each model and window we computed the residual (expected minus observed SNV count) and the standardized residual, defined as for Gnocchi (Results). *Global bias* is the squared difference between learned and true rates, averaged over windows. *Feature-specific bias* is the mean standardized residual as a function of feature value. The simulation shown in Figure 1 can be found [here](http://github.com/quinlan-lab/constraint-tools/blob/main/papers/neutral_models_are_biased/9.regression/simulation.4.2.1.interplay-of-SNR-and-bias.ipynb).

### Provenance of constraint scores

Gnocchi scores for a set of nonoverlapping 1000bp windows (“Chen windows”) were obtained from [here](https://www.nature.com/articles/s41586-023-06045-0#Sec28) (Supplementary Data #2). Windows on the X and Y chromosomes were omitted. Drs Nurdan Kuru and Adam Siepel ran ExtRaINSIGHT (Dukler et al. 2022) on the remaining Chen windows in order to compute λ_*S*_, excluding Chen windows that did not pass the ExtRaINSIGHT filters (private communication). A set of overlapping 500bp windows (“Halldorsson windows”) with Depletion Rank scores (Halldorsson et al. 2022) was obtained from [here](https://static-content.springer.com/esm/art%3A10.1038%2Fs41586-022-04965-x/MediaObjects/41586_2022_4965_MOESM3_ESM.gz). A set of overlapping 550bp windows with CDTS scores (“CDTS windows”) (di Iulio et al. 2018) was obtained by private communication with Dr Amalio Telenti (Vir Biotechnology), and their coordinates were lifted over from hg19 to hg38.

### Assignment of genomic feature values to genomic windows

We computed GC content in 1kb intervals centered on the Chen, Halldorsson and CDTS windows using “bedtools nuc”, e.g., as described [here](https://github.com/quinlan-lab/constraint-tools/blob/main/experiments/germline-model/chen-et-al-2022/compute-GC-content-given-window-size-based-on-Chen-windows.sh) and [here](https://github.com/quinlan-lab/constraint-tools/blob/main/experiments/germline-model/chen-et-al-2022/get-windows-from-chen.sh). We obtained BGS values computed by (Murphy et al. 2023) from [here](https://github.com/sellalab/HumanLinkedSelectionMaps/raw/master/Bmaps/CADD_bestfit.tar.gz), which we converted to bed format and lifted over from hg19 to hg38 (see [here](https://github.com/quinlan-lab/constraint-tools/blob/main/download-process-data/background-selection/download-B-maps.sh)). We obtained the gBGC coefficients on a set of genomic intervals for the European population computed by (Glémin et al. 2015) from [here](https://genome.cshlp.org/content/25/8/1215/suppl/DC1), and lifted over the genomic coordinates from hg18 to hg38, as described [here](https://github.com/quinlan-lab/constraint-tools/blob/main/download-process-data/download-gBGC-coefficient.sh). We assigned BGS values and gBGC coefficients (B_M1star.EUR) to the Chen, Halldorsson and CDTS windows by intersecting them with the bed files of BGS and gBGC, e.g., as shown [here](https://github.com/quinlan-lab/constraint-tools/blob/main/experiments/germline-model/chen-et-al-2022/Chen_models_SNV_counts_best_for_most_frequent_windows.1.1.ipynb) and [here](https://github.com/quinlan-lab/constraint-tools/blob/main/experiments/germline-model/chen-et-al-2022/Chen_models_SNV_counts_best_for_most_frequent_windows.1.2.ipynb).

### Construction of the window sets to assess model bias, and the lax truth sets to assess constraint-prediction performance

We filtered out untrustworthy windows, defined as windows that overlap any of the following intervals: [gaps in the hg38 genome assembly](https://github.com/quinlan-lab/constraint-tools/blob/main/download-process-data/download-gaps-grch38.sh), [Encode “exclude regions”](https://github.com/quinlan-lab/constraint-tools/blob/main/download-process-data/download-encode-exclude-regions-grch38.sh) (Amemiya et al. 2019), and [regions with insufficient read coverage in Gnomad version 3](https://github.com/quinlan-lab/constraint-tools/blob/main/download-process-data/gnomad/v3/coverage/download-gnomad-coverage.sh) (Chen et al. 2024). Exons were computed as shown [here](https://github.com/quinlan-lab/constraint-tools/blob/main/download-process-data/download-gaps-grch38.sh) and merged using “bedtools merge”. Noncoding windows were defined to be Chen, Halldorsson and CDTS windows that don’t significantly overlap merged exons. Of the noncoding windows, those that don’t significantly overlap Genehancer enhancers (Fishilevich et al. 2017) were labeled “neutral”, and were used to assess model bias. Noncoding windows that do significantly overlap Genehancer enhancers were labeled “constrained”. Together the “neutral” and “constrained” windows comprised the lax truth set used to assess the performance of each of the four constraint metrics.

### Modeling the mapping from genomic features to constraint scores

To compare constraint scores reported on different scales, we replaced each score’s value in each window by its rank among the analysed windows, divided by the number of windows. Ranks are therefore uniformly distributed between 0 and 1, with mean 0.5. In Figure 2A–C, the dark-grey line is the mean rank in each feature bin. Its slope is the marginal effect of a feature: its direct effect plus the effects of the features correlated with it. The direct effects are the coefficients of a linear regression of rank on all three standardized features jointly (light-grey line; Table 1). The co-variation of features with one another allows the linear regression (light-grey line) to track the observed nonlinear behavior (dark-grey line), e.g., in Figure 2B. The code to compute the linear regressions of the constraint scores shown in the heat maps, including the regression coefficients and the proportion of variance explained, reported in the tables, can be found [here](https://github.com/quinlan-lab/constraint-tools/blob/main/papers/neutral_models_are_biased/9.regression/experiment.1.ipynb).

### The expected distribution of standardized residuals on neutral windows

For Figure 2D, we selected windows whose GC content lies within 0.1 standard deviations of the mean. The expected curve is a standard normal density scaled by the histogram’s bin width and the number of windows. A standard normal is the distribution of Gnocchi expected under a simplified Chen model, as derived [here](https://github.com/quinlan-lab/constraint-tools/blob/main/papers/neutral_models_are_biased/10.residuals-are-over-dispersed.ipynb).

### Computation of window residuals under the Chen model

Each window has a residual, *R*, equal to the difference between the expected SNV count under the Chen model of neutrality, *μ*, and the observed SNV count, *S*. We computed *S* from Gnomad Version 3, using the bash command “constraint-tools predict-germline-model-Nonly --windows ${chen_windows} REQUIRED_ARGUMENTS”, where REQUIRED_ARGUMENTS are specified [here](https://github.com/quinlan-lab/constraint-tools/tree/main?tab=readme-ov-file#usage). We then used *S*, in conjunction with Gnocchi,*G*, to compute *μ*using the following algorithm:

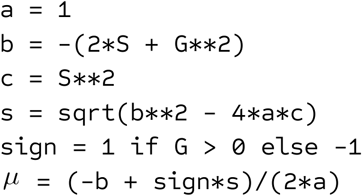

Finally:

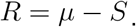

### Simulation of constraint calling (Figure 3)

Figures 1 and 3 come from a single simulation. Before fitting any model, we labelled a random 30% (300,000) of the 1,000,000 windows “constrained” and reduced each one’s SNV count by Δ = 15; both values were chosen to be representative of real data (Supporting Figure 2). The remaining windows were labelled “neutral”. The three models of Figure 1 were fit to the neutral windows only, as a model of neutral diversity should be, and Figure 1 shows only those windows. We then computed each window’s residual (expected minus observed SNV count) under each fitted model, for neutral and constrained windows alike. Because the constrained windows were excluded from the fits, the learned rates do not absorb the depletion, so a constrained window has a residual larger by Δ, on average, than a neutral window with the same feature value. For each model, we predicted a window to be constrained when its residual exceeded a threshold. Precision is the fraction of called windows that are labelled constrained, and recall is the fraction of constrained windows that are called. Lowering the threshold calls more windows, raising recall and typically lowering precision; varying it over the range of residuals traces one precision–recall curve per model. We did this separately for the windows in six strips of feature value: narrow (width 0.1) and wide (width 1) strips centred on 0, 1 and 2. Local bias is the global bias of Figure 1 computed over the windows in one strip. SNR was computed from the quadratic model’s residuals in the narrow strips (see “Computation of SNR” below). Supporting Figure 3 repeats the analysis using standardized residuals. Supporting Figure 4 repeats it with a depletion proportional to the feature value, in which case SNR rises, rather than falls, toward the right tail of the feature distribution. The code can be found [here](https://github.com/quinlan-lab/constraint-tools/blob/main/papers/neutral_models_are_biased/9.regression/simulation.4.2.1.interplay-of-SNR-and-bias.ipynb).

### Computation of the PR curve for the random classifier

Given a truth set of size *n*, let *π* represent the proportion of windows that are constrained. Let *p* represent the probability that a random classifier predicts that a given window is constrained.

Then

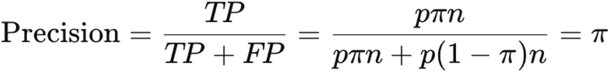

and

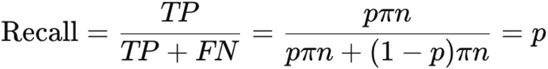

where TP (average number of True Positives) is the average number of windows that are labeled “constrained” in the truth set *and* are predicted to be constrained by the classifier. Similar definitions exist for FP (average number of False Positives) and FN (average number of False Negatives). The formulae for Precision and Recall describe a horizontal line on the Precision-Recall curve (e.g., the dashed line in Figure 4A, traced out as the parameter *p* is varied between 0 and 1) and represents a baseline level of performance that any meaningful constraint predictor ought to exceed.

### Computation of SNR

The signal-to-noise ratio (SNR) is defined as

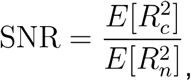

where *R*_*C*_ is the random residual of a constrained window,*R*_*n*_ is the random residual of a neutral window, and *E* [·] represents the expectation operator. The numerator is the amplitude of the meaningful signal we are trying to recover (a dip in SNV count due to purifying selection); the denominator is the amplitude of the unwanted background noise we must contend with (unavoidable Poisson fluctuations in SNV count). Squaring the relation

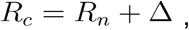

where Δ is the reduction in SNV count induced by purifying selection, we arrive at:

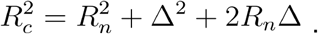

Applying the expectation operator, *E* [·], we get:

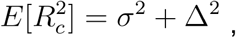

where we used the facts that Δ is constant (if Δ is feature-dependent, then specialize this analysis to a particular bin of that feature),*E*[*R*_*n*_]=0, and 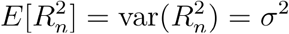.

Substituting these expressions for 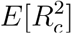 and 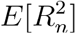 into the definition of SNR, we get

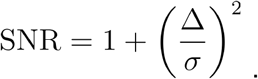

When Δ ≪ *σ*, SNR is approximately 1, indicating that the amplitudes of the signal and noise are approximately the same, making it impossible to recover the signal. This is reflected in a flat Precision-Recall (PR) curve (e.g., the dashed line in Figure 4A), which represents a baseline level of performance that any meaningful constraint predictor ought to exceed. We used the SNR formula to compute the numerical values of SNR reported in Figure 3 (and in Supporting Figure 4, where Δ is feature-dependent).

### Why PR curves for all models overlap when there is little variability in the mock genomic feature (**Figure 3G-I**)

Consider a set of simulated windows each with approximately the same value of the mock genomic feature. Under a given model, the predicted SNV count will be approximately the same for all these windows, because these models depend upon only that genomic feature. Now subset the windows into those that are constrained, and those that are not (neutral). The residual distributions for the two classes of windows (constrained and neutral) are just the observed SNV count distributions for the two classes, offset by a constant–the common predicted SNV count. Therefore the overlap of the residual distributions of the two classes is approximately the same for all models, and so all models perform approximately equally.

### Construction of the stringent truth set to assess performance of constraint scores

Chen et al (Chen et al. 2024) obtained a set of highly constrained genes (ClinGen haploinsufficient genes, MGI essential genes, OMIM autosomal dominant genes, LOEUF first-decile genes), linked enhancers to each gene using correlated patterns of histone modifications and gene expression, and assigned the most constrained enhancer to each gene, yielding a set of enhancers that we downloaded from [here](https://www.nature.com/articles/s41586-023-06045-0#Sec28) (Supplementary Data #6). We assigned genomic feature values, and constraint scores, to each of these “essential” enhancers similarly to how we did the assignments for the Chen, Halldorsson and CDTS windows. Finally, we constructed the stringent truth set by combining these “constrained” windows with an equal number of “neutral” windows sampled from the Chen lax truth set. See [here](https://github.com/quinlan-lab/constraint-tools/blob/main/papers/neutral_models_are_biased/8.labeled-enhancers/main.2.ipynb).

### Measuring constraint-prediction performance (Figure 4 and Supporting Figure 5)

For each constraint metric, we called a window constrained when its score passed a threshold, oriented so that higher values indicate constraint (we used the complements of Depletion Rank and of CDTS’s percentile rank), and computed precision and recall against the lax or stringent truth sets. Varying the threshold traces a precision–recall curve. A random classifier has precision equal to the fraction π of windows labelled constrained at every recall (“Computation of the PR curve for the random classifier”), so its area under the curve (auPRC) is π. We therefore report the normalized auPRC, auPRC/π, which is 1 for a random classifier. For a metric better than random, normalized auPRC still depends on π, since its maximum, 1/π, falls as π rises. We therefore compared GC-content bins at a common π: for each metric, we downsampled the constrained windows in every bin so that each bin’s ratio of constrained to neutral windows equaled the smallest such ratio among the bins (Figure 4A, B). Bins with fewer than 4,000 windows after downsampling are not shown. To compare the lax and stringent truth sets (Figure 4C, D), we drew a random subsample of the lax truth set of the same size as the stringent truth set, and generated 1,000 bootstrap samples of each. In each bootstrap sample, we kept the windows with GC content in (0.2, 0.375] or (0.4, 0.7] and downsampled the constrained windows so that the two bins had a common π, as above. We then computed normalized auPRC over the windows of both bins (C), and the auPRC of the low-GC bin minus that of the high-GC bin, divided by the auPRC over both bins (D). We report the mean and standard deviation over bootstrap samples. Supporting Figure 5 repeats these analyses with windows binned by BGS and by gBGC. Code for (A, B) is [here](https://github.com/quinlan-lab/constraint-tools/blob/main/papers/neutral_models_are_biased/7.CDTS/main.2.ipynb) and for (C, D) is [here](https://github.com/quinlan-lab/constraint-tools/blob/main/papers/neutral_models_are_biased/11.compare-lax-with-stringent-truth-set.ipynb). Note that downsampling gives the bins compared within a figure a common π, but not the two truth sets, the four metrics, or different figures. We therefore repeated each of those comparisons with the area under the receiver operating characteristic curve (auROC), which does not depend on π, obtaining qualitatively similar results ([code](https://github.com/petermchale/gnomad_nc_constraint/blob/main/fig4_prevalence/main.2.prevalence.ipynb) for Figure 4A, B and [code](https://github.com/petermchale/gnomad_nc_constraint/blob/main/fig4_prevalence/11.compare-lax-with-stringent-truth-set.prevalence.ipynb) for Figure 4C, D and Supporting Figure 5E–H). Normalized auPRC should be compared only within a figure: Figure 4C and Supporting Figure 5E, F downsample by different features, resulting in different final values of π.

### Mathematical and computational dissection, and correction, of Gnocchi’s GC bias

We verified that we could reproduce Gnocchi using code and intermediate data files supplied by Chen et al ([link](https://github.com/petermchale/gnomad_nc_constraint/tree/main/preconditions)). The mathematical derivations and code we used to produce Fig 5, Supporting Figure 7, and Supporting Figure 8 can be found [here](https://github.com/petermchale/gnomad_nc_constraint/blob/main/fig5/fig5.ipynb). We used Claude Code in a significant fraction of these analyses, and to help frame but not write some of the corresponding arguments in the manuscript.

## Acknowledgements

We graciously thank Nurdan Kuru and Adam Siepel for sharing their ExtRaINSIGHT data. We are grateful to Andrew Kern for expert feedback that significantly improved on an early draft of the manuscript.

## References

Adrion, Jeffrey R., Christopher B. Cole, Noah Dukler, et al. 2020. “A Community-Maintained Standard Library of Population Genetic Models.” eLife 9 (e54967). 10.7554/eLife.54967.

Amemiya, Haley M., Anshul Kundaje, and Alan P. Boyle. 2019. “The ENCODE Blacklist: Identification of Problematic Regions of the Genome.” Scientific Reports 9 (1): 9354.

Amos, W., and J. I. Hoffman. 2010. “Evidence That Two Main Bottleneck Events Shaped Modern Human Genetic Diversity.” Proceedings. Biological Sciences 277 (1678): 131–137.

Arndt, Peter F., Terence Hwa, and Dmitri A. Petrov. 2005. “Substantial Regional Variation in Substitution Rates in the Human Genome: Importance of GC Content, Gene Density, and Telomere-Specific Effects.” Journal of Molecular Evolution 60 (6): 748–763.

Bajwa, Ayesha, Ruchir Rastogi, Pooja Kathail, Richard W. Shuai, and Nilah Ioannidis. 2024. “Characterizing Uncertainty in Predictions of Genomic Sequence-to-Activity Models.” Machine Learning in Computational Biology, March 15, 279–297.

Benegas, Gonzalo, Carlos Albors, Alan J. Aw, Chengzhong Ye, and Yun S. Song. 2024. “GPN-MSA: An Alignment-Based DNA Language Model for Genome-Wide Variant Effect Prediction.” bioRxiv.org: The Preprint Server for Biology, ahead of print, April 6. 10.1101/2023.10.10.561776.

Bishop, Christopher M. 2006. Pattern Recognition and Machine Learning. 1 (August): 740.

Boffelli, Dario, Jon McAuliffe, Dmitriy Ovcharenko, et al. 2003. “Phylogenetic Shadowing of Primate Sequences to Find Functional Regions of the Human Genome.” Science (New York, N.Y.) 299 (5611): 1391–1394.

Bulmer, M. 1986. “Neighboring Base Effects on Substitution Rates in Pseudogenes.” Molecular Biology and Evolution 3 (4): 322–329.

Chandra, Sheel, and Ziyue Gao. 2025. “Sequence Context and Methylation Interact to Shape Germline Mutation Rate Variation at CpG Sites.” In bioRxiv. November 13. 10.1101/2025.11.13.688199.

Charlesworth, Brian. 2013. “Background Selection 20 Years on: The Wilhelmine E. Key 2012 Invitational Lecture.” The Journal of Heredity 104 (2): 161–171.

Chen, Siwei, Laurent C. Francioli, Julia K. Goodrich, et al. 2024. “A Genomic Mutational Constraint Map Using Variation in 76,156 Human Genomes.” Nature 625 (7993): 92–100.

Cooper, Gregory M., Eric A. Stone, George Asimenos, et al. 2005. “Distribution and Intensity of Constraint in Mammalian Genomic Sequence.” Genome Research 15 (7): 901–913.

Davydov, Eugene V., David L. Goode, Marina Sirota, Gregory M. Cooper, Arend Sidow, and Serafim Batzoglou. 2010. “Identifying a High Fraction of the Human Genome to Be under Selective Constraint Using GERP++.” PLoS Computational Biology 6 (12): e1001025.

Dukler, Noah, Mehreen R. Mughal, Ritika Ramani, Yi-Fei Huang, and Adam Siepel. 2022. “Extreme Purifying Selection against Point Mutations in the Human Genome.” Nature Communications 13 (1): 4312.

Duret, Laurent, and Nicolas Galtier. 2009. “Biased Gene Conversion and the Evolution of Mammalian Genomic Landscapes.” Annual Review of Genomics and Human Genetics 10: 285–311.

Elango, Navin, Seong-Ho Kim, Eric Vigoda, and Soojin V. Yi. 2008. “Mutations of Different Molecular Origins Exhibit Contrasting Patterns of Regional Substitution Rate Variation.” PLoS Computational Biology 4 (2): e1000015.

Fang, Yiyuan, Shuyi Deng, and Cai Li. 2022. “A Generalizable Deep Learning Framework for Inferring Fine-Scale Germline Mutation Rate Maps.” Nature Machine Intelligence 4 (12): 1209–1223.

Fishilevich, Simon, Ron Nudel, Noa Rappaport, et al. 2017. “GeneHancer: Genome-Wide Integration of Enhancers and Target Genes in GeneCards.” Database: The Journal of Biological Databases and Curation 2017 (January): bax028.

Friedman, Jerome H. 2001. “Greedy Function Approximation: A Gradient Boosting Machine.” Annals of Statistics 29 (5): 1189–1232.

Fryxell, Karl J., and Won-Jong Moon. 2005. “CpG Mutation Rates in the Human Genome Are Highly Dependent on Local GC Content.” Molecular Biology and Evolution 22 (3): 650–658.

Glémin, Sylvain, Peter F. Arndt, Philipp W. Messer, Dmitri Petrov, Nicolas Galtier, and Laurent Duret. 2015. “Quantification of GC-Biased Gene Conversion in the Human Genome.” Genome Research 25 (8): 1215–1228.

Gower, Graham, Nathaniel S. Pope, Murillo F. Rodrigues, et al. 2025. “Accessible, Realistic Genome Simulation with Selection Using Stdpopsim.” In bioRxivorg. August 12. 10.1101/2025.03.23.644823.

Halldorsson, Bjarni V., Hannes P. Eggertsson, Kristjan H. S. Moore, et al. 2022. “The Sequences of 150,119 Genomes in the UK Biobank.” Nature 607 (7920): 732–740.

Huber, Christian D., Bernard Y. Kim, and Kirk E. Lohmueller. 2020. “Population Genetic Models of GERP Scores Suggest Pervasive Turnover of Constrained Sites across Mammalian Evolution.” PLoS Genetics 16 (5): e1008827.

Iulio, Julia di, Istvan Bartha, Emily H. M. Wong, et al. 2018. “The Human Noncoding Genome Defined by Genetic Diversity.” Nature Genetics 50 (3): 333–337.

Kathail, Pooja, Richard W. Shuai, Ryan Chung, Chun Jimmie Ye, Gabriel B. Loeb, and Nilah M. Ioannidis. 2024a. “Current Genomic Deep Learning Models Display Decreased Performance in Cell Type Specific Accessible Regions.” bioRxiv.org: The Preprint Server for Biology, July 10, 2024.07.05.602265.

Kathail, Pooja, Richard W. Shuai, Ryan Chung, Chun Jimmie Ye, Gabriel B. Loeb, and Nilah M. Ioannidis. 2024b. “Current Genomic Deep Learning Models Display Decreased Performance in Cell Type-Specific Accessible Regions.” Genome Biology 25 (1): 202.

Lauterbur, M. Elise, Maria Izabel A. Cavassim, Ariella L. Gladstein, et al. 2023. “Expanding the Stdpopsim Species Catalog, and Lessons Learned for Realistic Genome Simulations.” May 23. 10.7554/elife.84874.

Liu, Li, Maxwell D. Sanderford, Ravi Patel, Pramod Chandrashekar, Greg Gibson, and Sudhir Kumar. 2019. “Biological Relevance of Computationally Predicted Pathogenicity of Noncoding Variants.” Nature Communications 10 (1): 330.

Lohmueller, Kirk E. 2014. “The Distribution of Deleterious Genetic Variation in Human Populations.” Current Opinion in Genetics & Development 29 (December): 139–146.

Luikart, G., F. W. Allendorf, J. M. Cornuet, and W. B. Sherwin. 1998. “Distortion of Allele Frequency Distributions Provides a Test for Recent Population Bottlenecks.” The Journal of Heredity 89 (3): 238–247.

Martini, Lorenzo, Roberta Bardini, Alessandro Savino, and Stefano Di Carlo. 2023. “GRAIGH: Gene Regulation Accessibility Integrating GeneHancer Database.” 2023 IEEE International Conference on Bioinformatics and Biomedicine (BIBM), December 5, 343–348.

McVicker, Graham, David Gordon, Colleen Davis, and Phil Green. 2009. “Widespread Genomic Signatures of Natural Selection in Hominid Evolution.” PLoS Genetics 5 (5): e1000471.

Meader, Stephen, Chris P. Ponting, and Gerton Lunter. 2010. “Massive Turnover of Functional Sequence in Human and Other Mammalian Genomes.” Genome Research 20 (10): 1335–1343.

Morcos, Faruck, Andrea Pagnani, Bryan Lunt, et al. 2011. “Direct-Coupling Analysis of Residue Coevolution Captures Native Contacts across Many Protein Families.” Proceedings of the National Academy of Sciences of the United States of America 108 (49): E1293–301.

Mugal, Carina F., and Hans Ellegren. 2011. “Substitution Rate Variation at Human CpG Sites Correlates with Non-CpG Divergence, Methylation Level and GC Content.” Genome Biology 12 (6): R58.

Murphy, David A., Eyal Elyashiv, Guy Amster, and Guy Sella. 2023. “Broad-Scale Variation in Human Genetic Diversity Levels Is Predicted by Purifying Selection on Coding and Non-Coding Elements.” eLife 12 (June). 10.7554/eLife.76065.

Pollard, Katherine S., Melissa J. Hubisz, Kate R. Rosenbloom, and Adam Siepel. 2010. “Detection of Nonneutral Substitution Rates on Mammalian Phylogenies.” Genome Research 20 (1): 110–121.

Ponting, Chris P., and Ross C. Hardison. 2011. “What Fraction of the Human Genome Is Functional?” Genome Research 21 (11): 1769–1776.

Pouyet, Fanny, Simon Aeschbacher, Alexandre Thiéry, and Laurent Excoffier. 2018. “Background Selection and Biased Gene Conversion Affect More than 95% of the Human Genome and Bias Demographic Inferences.” eLife 7 (August). 10.7554/eLife.36317.

Rands, Chris M., Stephen Meader, Chris P. Ponting, and Gerton Lunter. 2014. “8.2% of the Human Genome Is Constrained: Variation in Rates of Turnover across Functional Element Classes in the Human Lineage.” PLoS Genetics 10 (7): e1004525.

Ratnakumar, Abhirami, Sylvain Mousset, Sylvain Glémin, et al. 2010. “Detecting Positive Selection within Genomes: The Problem of Biased Gene Conversion.” Philosophical Transactions of the Royal Society of London. Series B, Biological Sciences 365 (1552): 2571–2580.

Schraiber, Joshua G., Jeffrey P. Spence, and Michael D. Edge. 2024. “Estimation of Demography and Mutation Rates from One Million Haploid Genomes.” In Genomics, No. Biorxiv;2024.09.18.613708v1. BioRxiv, September 22. https://www.biorxiv.org/content/10.1101/2024.09.18.613708v1.full.pdf.

Schrider, Daniel R., and Andrew D. Kern. 2015. “Inferring Selective Constraint from Population Genomic Data Suggests Recent Regulatory Turnover in the Human Brain.” Genome Biology and Evolution 7 (12): 3511–3528.

Smith, J. M., and J. Haigh. 1974. “The Hitch-Hiking Effect of a Favourable Gene.” Genetical Research 23 (1): 23–35.

Smith, Nick G. C., Mikael Brandström, and Hans Ellegren. 2004. “Evidence for Turnover of Functional Noncoding DNA in Mammalian Genome Evolution.” Genomics 84 (5): 806–813.

Stephan, Wolfgang. 2019. “Selective Sweeps.” Genetics 211 (1): 5–13.

Sved, J., and A. Bird. 1990. “The Expected Equilibrium of the CpG Dinucleotide in Vertebrate Genomes under a Mutation Model.” Proceedings of the National Academy of Sciences of the United States of America 87 (12): 4692–4696.

Telis, Natalie, Robin Aguilar, and Kelley Harris. 2020. “Selection against Archaic Hominin Genetic Variation in Regulatory Regions.” Nature Ecology & Evolution 4 (11): 1558–1566.

Trevino, Alexandro E., Fabian Müller, Jimena Andersen, et al. 2021. “Chromatin and Gene-Regulatory Dynamics of the Developing Human Cerebral Cortex at Single-Cell Resolution.” Cell 184 (19): 5053–5069.e23.

Venkat, Aarti, Matthew W. Hahn, and Joseph W. Thornton. 2018. “Multinucleotide Mutations Cause False Inferences of Lineage-Specific Positive Selection.” Nature Ecology & Evolution 2 (8): 1280–1288.

Ward, Lucas D., and Manolis Kellis. 2012. “Evidence of Abundant Purifying Selection in Humans for Recently Acquired Regulatory Functions.” Science (New York, N.Y.) 337 (6102): 1675–1678.

Yap, Von Bing, Helen Lindsay, Simon Easteal, and Gavin Huttley. 2010. “Estimates of the Effect of Natural Selection on Protein-Coding Content.” Molecular Biology and Evolution 27 (3): 726–734.

