## Supplementary Information for "The performance of genetic-constraint metrics varies significantly across the human noncoding genome"

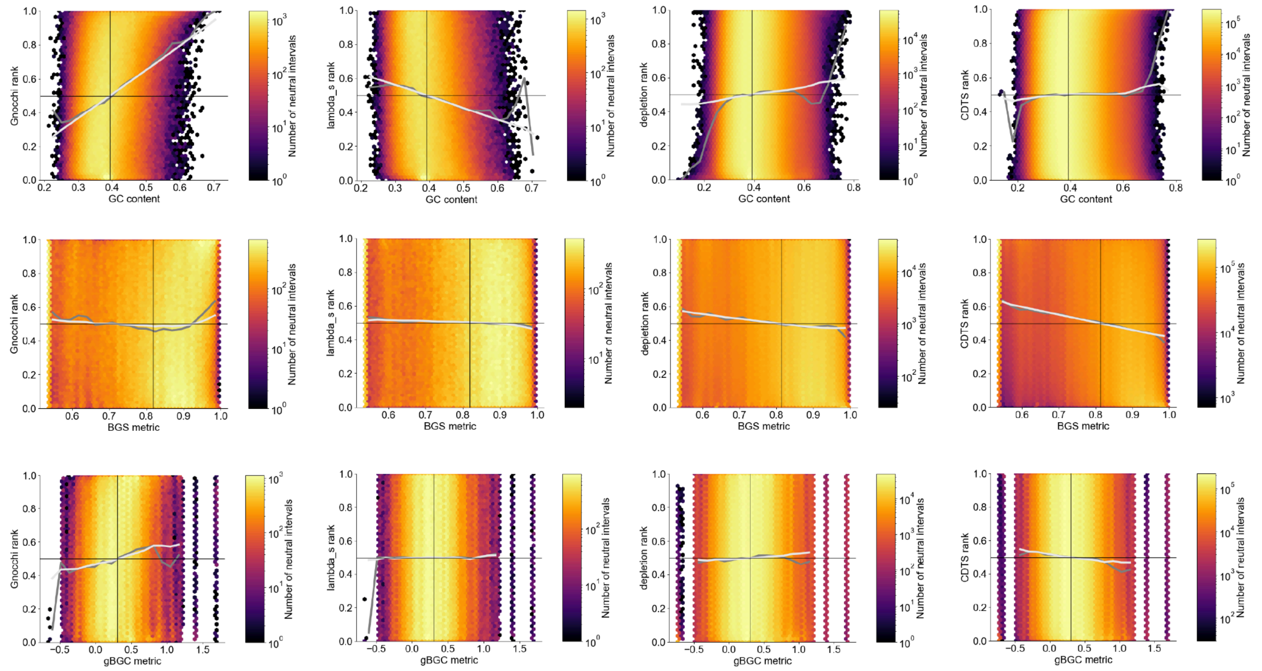


**Supporting Figure 1** Similarities and differences in the biases observed in four different constraint metrics (columns) as a function of three different features (rows). Ranks are standardized to lie in the unit interval.

A

| window overlaps enhancer | number of windows |
|---|---|
| False | 693270 |
| True | 309957 |

B

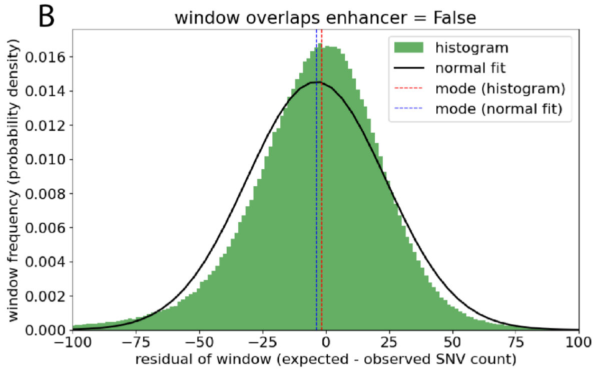


C

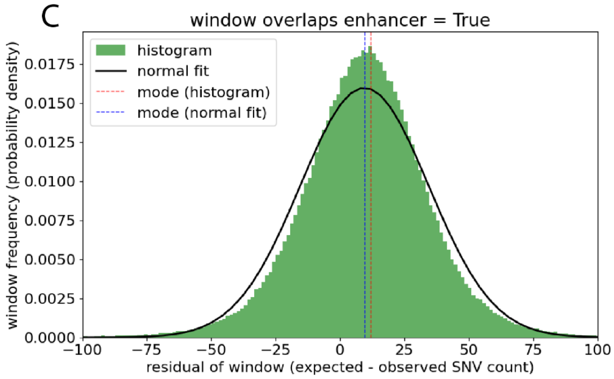


D

| | |
|---|---|
| **depletion (estimated using histogram)** | 13.676763 |
| **depletion (estimated using fit to histogram)** | 13.293118 |

**Supporting Figure 2** Using experimental data to estimate the imbalance in the number of neutral and constrained windows, and the depletion of SNV counts due to purifying selection in the constrained windows. (A) The number of noncoding Chen windows that overlap a Genehancer enhancer or not. The ratio of these numbers is an estimate of the imbalance between the numbers of neutral and constrained windows. (B and C) Each window has a residual, $R$, equal to the difference between the expected SNV count under the Chen model of neutrality, $\mu$, and the observed SNV count, $S$ (Methods). These plots show the distribution of those residuals over all noncoding Chen windows, broken down by whether the window is assumed to be constrained or not. In each case, the vertical red line indicates the mode of the distribution, estimated using the histogram; and the vertical blue line indicates the mode of the normal distribution that best fits the histogram. (D) The difference of the modes obtained from the two classes of windows (putatively constrained or not) is an estimate of the depletion of SNV count due to purifying selection, $\Delta$. Code to produce the figure can be found here. In light of these experimental results, we elected to use a class imbalance of 3:7 (constrained:unconstrained window count) and a depletion of $\Delta = 15$ SNVs due to (weak) purifying selection in the simulation shown in **Figure 3**.

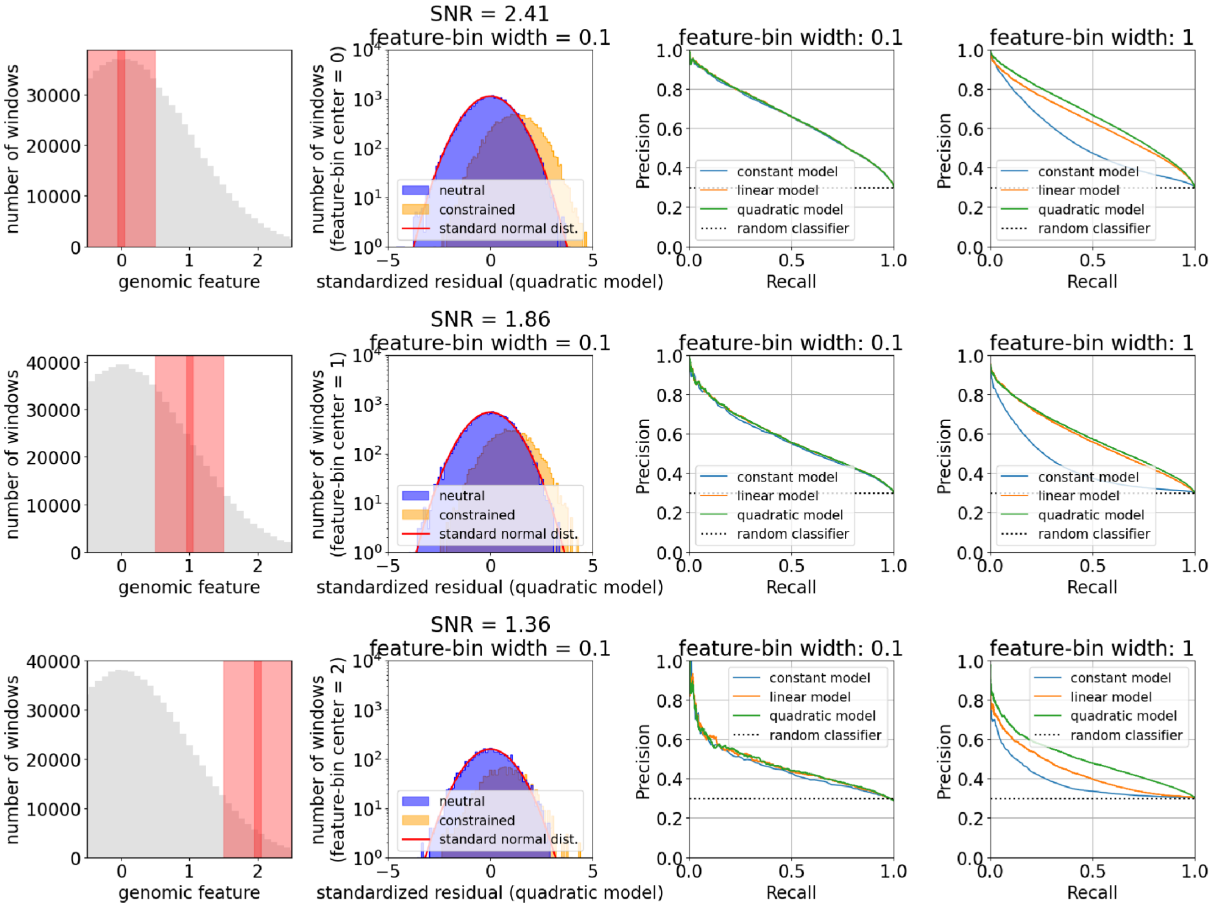


**Supporting Figure 3** Similar to **Figure 3**, but using standardized residuals, not raw residuals. The performance trends are identical to those seen in Figure 3. This is because the SNR computed using standardized residuals, $\mathrm{SNR} = E[G_c^2]/E[G_n^2]$, where $G = R/\sigma$, is identical to that computed using raw residuals, $\mathrm{SNR} = E[R_c^2]/E[R_n^2]$. Here, the subscripts $c$ and $n$ refer to "constrained" and "neutral", respectively.

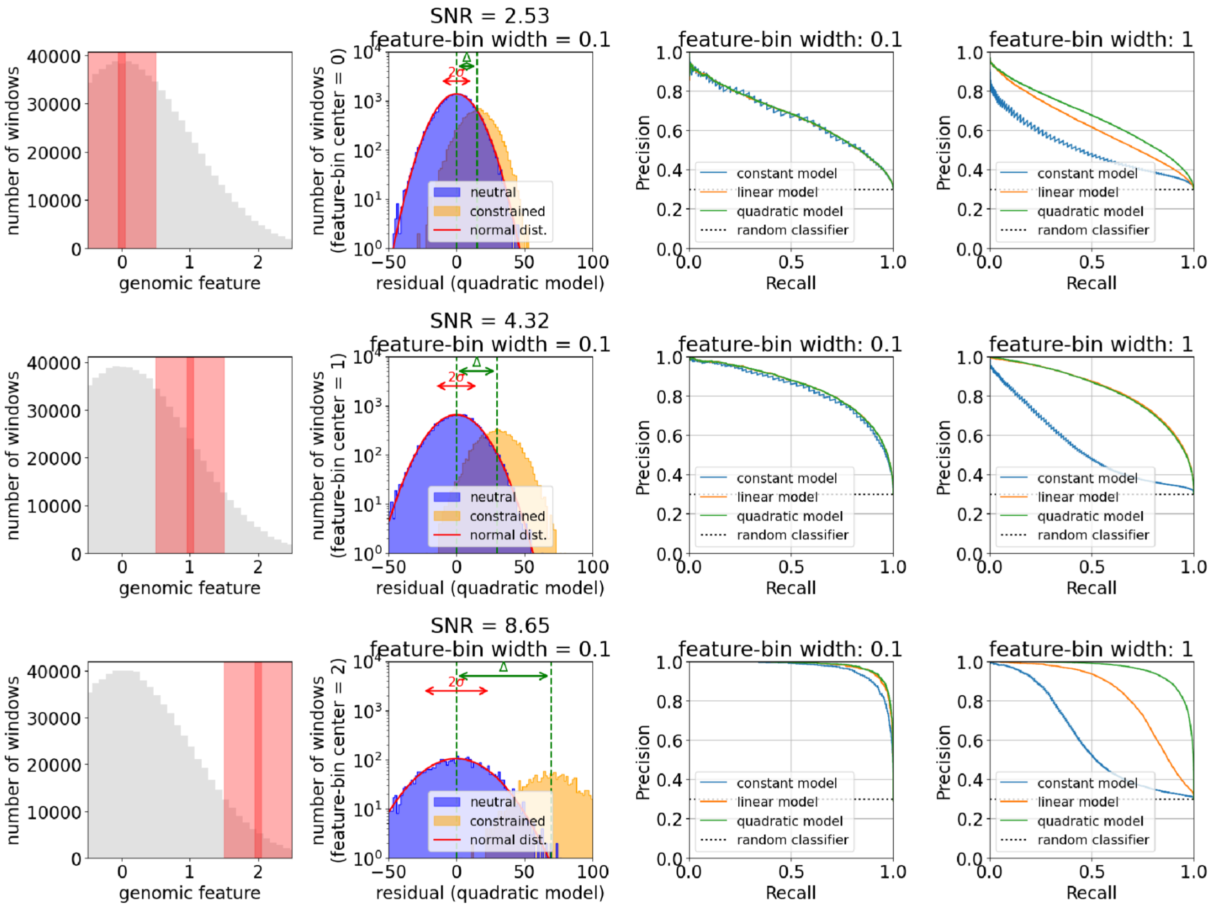


**Supporting Figure 4** SNR and model bias are both expected to significantly affect the ability of neutral models to call constraint. Similar to **Figure 3**, except that SNV counts are depleted by an amount $\Delta$ proportional to the feature value (instead of a feature-independent $\Delta$, as in **Figure 3**), to mimic scenarios where constrained windows tend to have larger feature values, e.g., enhancers are enriched in GC content. The simulation is here.

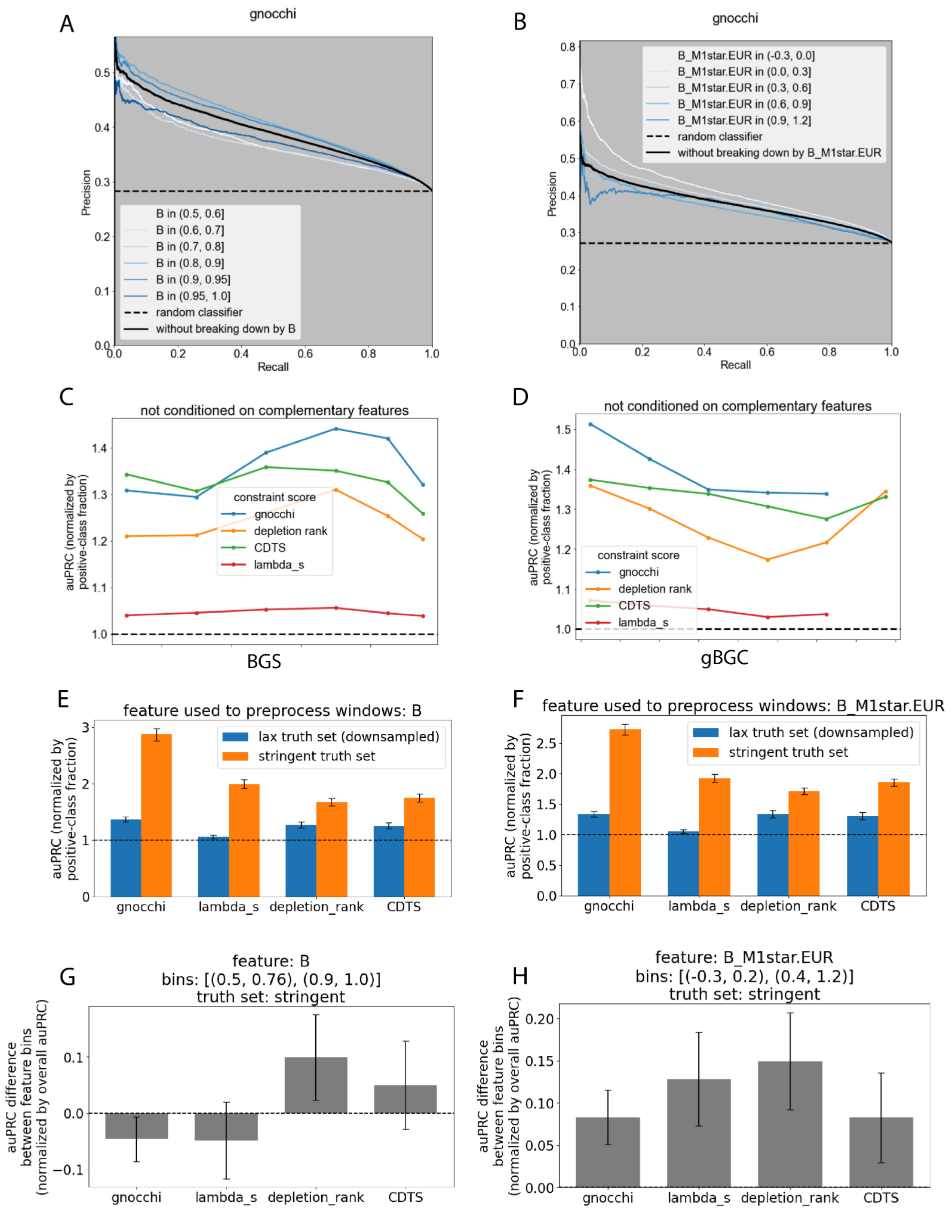


**Supporting Figure 5** The performance of neutral-diversity models at predicting constraint covaries with the nature of the genomic windows upon which performance is assessed. Here, we stratify by background selection (BGS) and GC-biased gene conversion (gBGC). (A, B) The performance of the Chen model at predicting constraint relative to a lax truth set, defined by whether windows overlap GeneHancer enhancers. (C, D) Performance (area under the precision-recall curve, auPRC, normalized by the fraction of examples that are positive) varies with BGS and gBGC in a largely similar way for all constraint metrics. (E, F) Performance increases on moving from the lax to the stringent truth set (defined by a set of essential genes), as expected, particularly for the Gnocchi and $\lambda_s$ metrics. (G, H) Performance varies with BGS and gBGC when assessed relative to the stringent truth set. The bar charts report the mean and standard deviation of bootstrapped differences between normalized auPRCs for two bins of each feature. Compare with panels C and D.

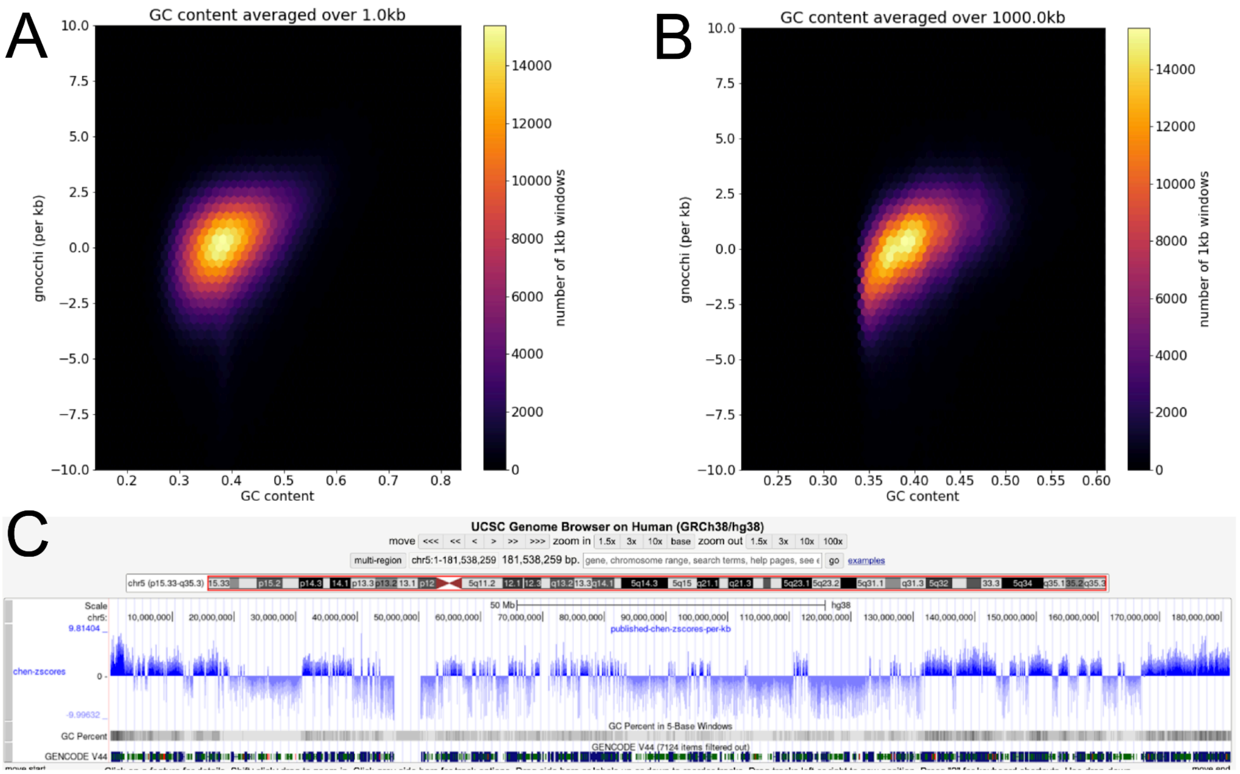


**Supporting Figure 6** The origin of Gnocchi's autocorrelation. (A, B) Gnocchi positively correlates with GC content at multiple scales. (C) The well-known autocorrelation of GC content ("isochores") therefore implies autocorrelation of Gnocchi.

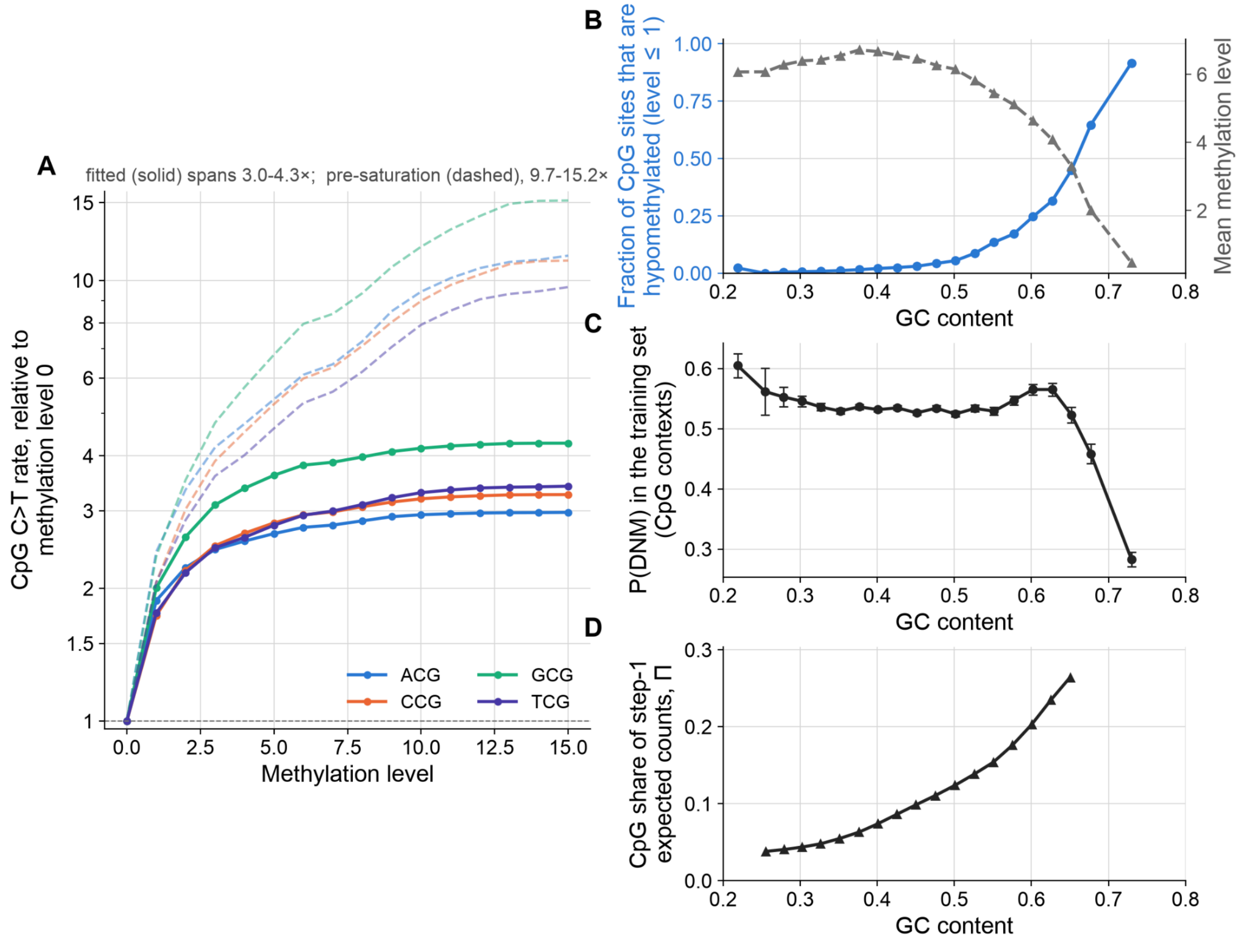


**Supporting Figure 7** The ablated, context-only model implicitly corrects for GC content. (A) Mutation rate at CpG sites increases with their DNA methylation level. Solid, the fitted C>T probability for each of the four CpG contexts against methylation level, each divided by its own value at level 0. Dashed, the same observable read in a 1,000-genome downsample and normalised the same way. The gap between them is saturation of the polymorphism probability at the larger cohort size, derived analytically in https://github.com/petermchale/gnomad_nc_constraint/blob/main/fig5/fig5.ipynb (B) High-GC-content CpGs (CpG islands) are hypomethylated. The fraction of CpG-context sites in the DNM-submodel's training set that are hypomethylated (methylation level <= 1; blue, left axis) and the mean methylation level (grey, right axis), per GC bin, covary with GC content. (C) The drop in methylation in CpG islands (panel B) causes their DNM rate to collapse (this panel). Empirical P(DNM) at CpG-context training sites per GC bin. (D) The increased density of CpG sites in CpG islands is reflected in the CpG share of expected SNV counts in 1kb windows under the context-only model (Figure 5B's decomposition). (A)-(C) are binned over the DNM training sites and (D) over 1kb windows which Gnocchi scored.

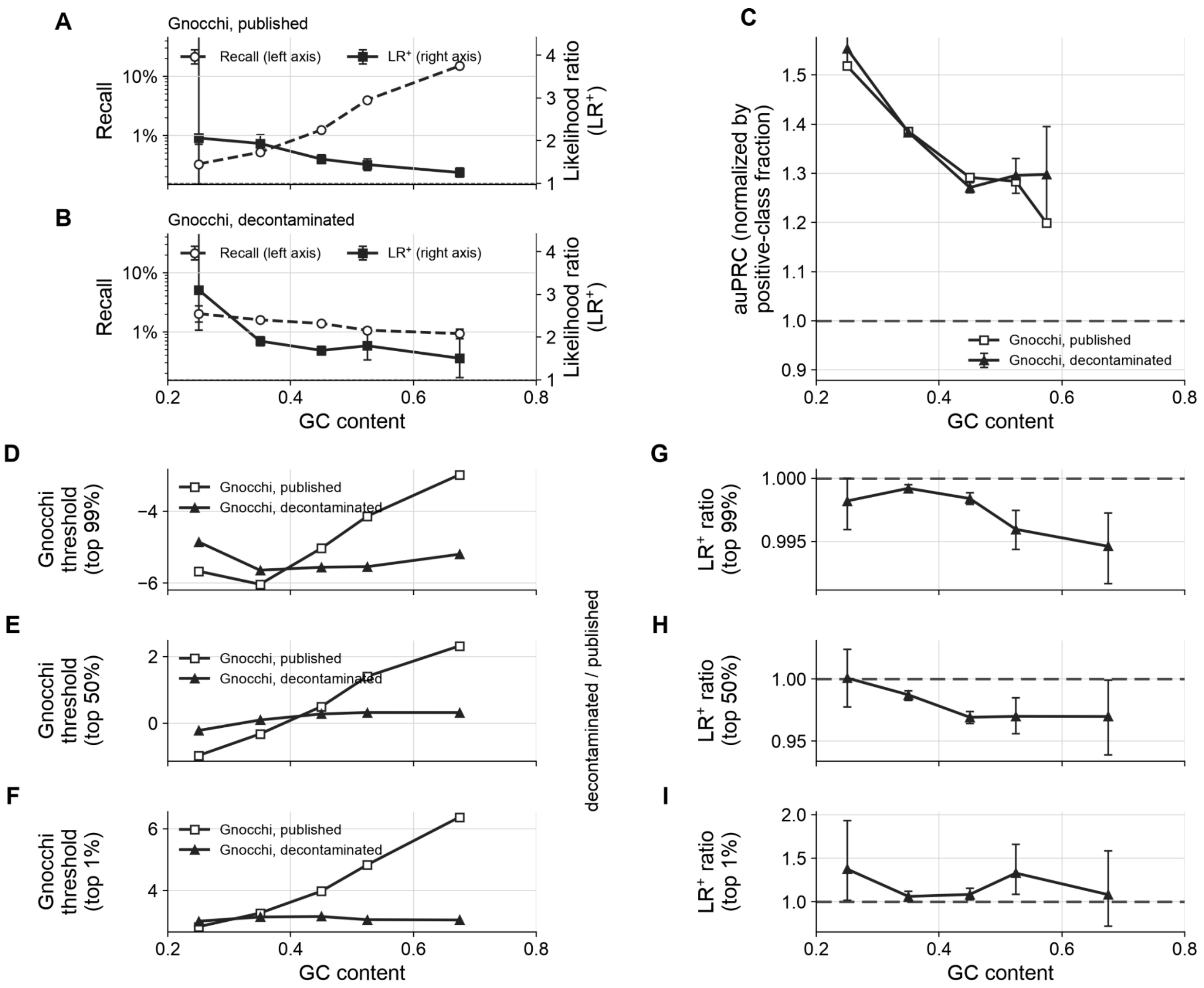


**Supporting Figure 8** Discovery performance of published and decontaminated Gnocchi on the lax truth set. (A, B) Decontaminating Gnocchi redistributes constraint discovery toward sequences where a call is worth more. We quantified the evidential weight of a call by the positive likelihood ratio LR+ = P(call | constrained) / P(call | not constrained), which conditions on the true class on both sides, and so is comparable between bins (which differ in prevalence). As for Figure 5F, the thresholds were chosen to match genomewide calling rates across scores, yielding, for this dataset, a threshold of 4 for published Gnocchi and a threshold of 3.14 for decontaminated Gnocchi. (C) Averaged over all Gnocchi thresholds, decontamination has no significant effect on discovery. Discriminative power falls with GC content regardless of whether the score is biased or not, implying that SNR, not bias, drives the declining performance of published Gnocchi as a function of GC content. As for Figure 4A, we downsampled the positive class to give every bin the same constraint prevalence. (D-I) When the discovery budget is fixed, however, the performance of published and decontaminated Gnocchi can be seen to diverge. Each row shows the threshold each score would need in order to call the indicated top percentage of every GC bin (left column), and the ratio of the corresponding evidential weights of calls made using those thresholds (right column). The indicated call-rate percentages correspond to the left tail, the median and the right tail of each GC-specific score distribution.

The smallest divergence occurs in the left tail (G) but neither score is particularly discriminative there anyway. (It's a mathematical fact that LR+ approaches 1, independent of score, as the corresponding threshold approaches negative infinity). In contrast, the largest divergence in evidential weights is seen in the right tail of the score distributions—the operating point of most applications (c.f., panel B for typical values of LR+ at this operating point). Here we find that decontamination moderately increases the evidential weight of calls (panels F and I).
